# Evidence for negative selection against somatic mutations in fibroblasts with high N-ethyl-N-nitrosourea-induced mutation load

**DOI:** 10.1101/2024.04.07.588286

**Authors:** Ronald Cutler, Johanna Heid, Shixiang Sun, Moonsook Lee, Alexander Y. Maslov, Lei Zhang, Simone Sidoli, Xiao Dong, Jan Vijg

## Abstract

Advanced sequencing has revealed that thousands of mutations accumulate with age in most human tissues. While some can clonally expand and cause disease, the maximum mutation load a cell can tolerate without functional decline is unknown. We addressed this by repeatedly treating proliferating human primary fibroblasts with a low dose of the mutagen N-ethyl-N-nitrosourea and quantifying somatic mutation burden using single-cell whole-genome sequencing. Mutation load increased linearly to ∼56,000 single-nucleotide variants per cell with only a slight reduction in growth rate. Analysis showed negative selection against potentially deleterious mutations in coding and non-coding regions, including sequences tied to pathways essential for cell growth and identity. This selective removal of harmful variants likely enables cells to maintain growth functions despite extreme mutation burden. Because most adult tissues are non-dividing and cannot benefit from negative selection based on growth, mutations that accumulate during aging may have pronounced functional consequences.

## Introduction

Aging is a complex process of continuous decline in cellular and tissue function that is accompanied with increased disease risk(*1*). DNA damage is thought to play a central role in this process and may be a universal cause of aging(*2*). Among the major molecular consequences of DNA damage during aging are somatic DNA mutations. Somatic mutations are the result of errors during repair or replication of a damaged DNA template and can only be purged via the death of the cell or organism. The accumulation of somatic mutations has since long been implicated as a major cause of aging(*3, 4*), which is strongly supported by the age-related exponential increase of cancer, a disease known to be caused by mutations(*5*). Somatic mutations are difficult to analyze because of their random nature, with very low variant allele frequencies indistinguishable from technical artifacts by traditional bulk-sequencing methods. With the recent emergence of single-cell and single-molecule assays, this situation has changed and we are now able to quantitatively analyze most small mutations, i.e. base substitutions and small insertions and deletions, at high accuracy. Application of such methods has provided ample evidence that somatic mutation rate is far higher than the germline mutation rate and that thousands of somatic mutations accumulate with age in most, if not all, human tissues(*6–10*), including postmitotic tissues such as the brain and heart(*11, 12*). In addition, evidence has emerged that postzygotic, somatic mutations are a cause of a wide variety of human diseases other than cancer(*13, 14*). Like cancer, many of these age-related effects may be due to clonally amplified mutations(*15–17*). Indeed, in blood, somatic mutations lead to clonal hematopoiesis, which has been associated with increased risk of age-related diseases, such as cardiovascular disease, as well as overall mortality(*18*).

What thus far remains unknown is the possible functional impact, collectively, of thousands of random mutations accumulating in a non-cancerous cell, for example, by adversely affecting gene-coding or gene-regulatory sequences relevant for the given cell type. If so, one would expect a limit to the burden of random mutations a cell can tolerate or selection against mutations in important functional regions. Here, we directly tested this hypothesis by treating actively proliferating human primary fibroblasts with a low, repeated dose of N-ethyl-N-nitrosourea (ENU), a powerful mutagen that induces almost exclusively single-nucleotide variants (SNVs). Our results show that 9 cycles of ENU treatment over the course of ∼24 population doublings linearly increased mutation frequency up to ∼56,000 SNVs per cell, with only subtle effects on cell growth and death rates. We show that this tolerance for somatic mutations could be explained by negative selection against potentially deleterious SNVs in coding and non-coding genomic loci as well as selection against SNVs across the functional genomic loci associated with cell growth and cell identity. These results suggest that accumulating somatic mutations may have adverse effects on cellular function when negative selection is not possible, as is the case in most cells and tissues in adult organisms, which are not mitotically active(*19*).

## Results

### Repeated Mutagen Treatment of Normal Cells Only Slightly Affects Cell Growth and Death

To determine the impact of increased somatic mutation burden on primary human cells *in vitro*, we treated low-passage fetal lung fibroblasts (IMR-90) multiple times with a sublethal dose of 50 µg/mL of ENU. Each treatment was followed by a recovery period of 7 days, after which 1 million cells (determined by cell counting) were replated and treated again. This process was repeated for 9 cycles (Figure 1A; Methods). To monitor cell growth and toxicity, we measured cell number and apoptosis rate at 3 and 7 days after each treatment to assess both short- and long-term effects of the treatment (Methods). We measured cell growth by calculating population doublings (PDs), and found that repeated treatment with ENU gradually decreased cell growth rate over the course of the experiment, as at cycle 9 there were a total of 28.25±1.87 (mean ± standard deviation) PDs for control cells compared to 24.1±1.34 PDs for ENU-treated cells (Figure 1B). The effect of ENU treatment on cell growth was more pronounced at the 3 day time point (Figure 1C) compared to the 7 day time point (Figure 1D), which mostly explains the decrease in overall growth rate (Figure 1B). A larger effect on growth rate at the 3 day timepoint was expected due to the DNA damage response that is known to occur early and then subsides over time. To identify the contribution of cell death to the slight decrease in growth rate due to mutagen exposure, we measured markers of early and late apoptosis at the 3 and 7 day time points. The results showed a slight, but consistent increase in early and late apoptosis in the ENU-treated group as compared to the control cells at both time points (Figures 1E-F and 1G-H).

**Figure 1.**
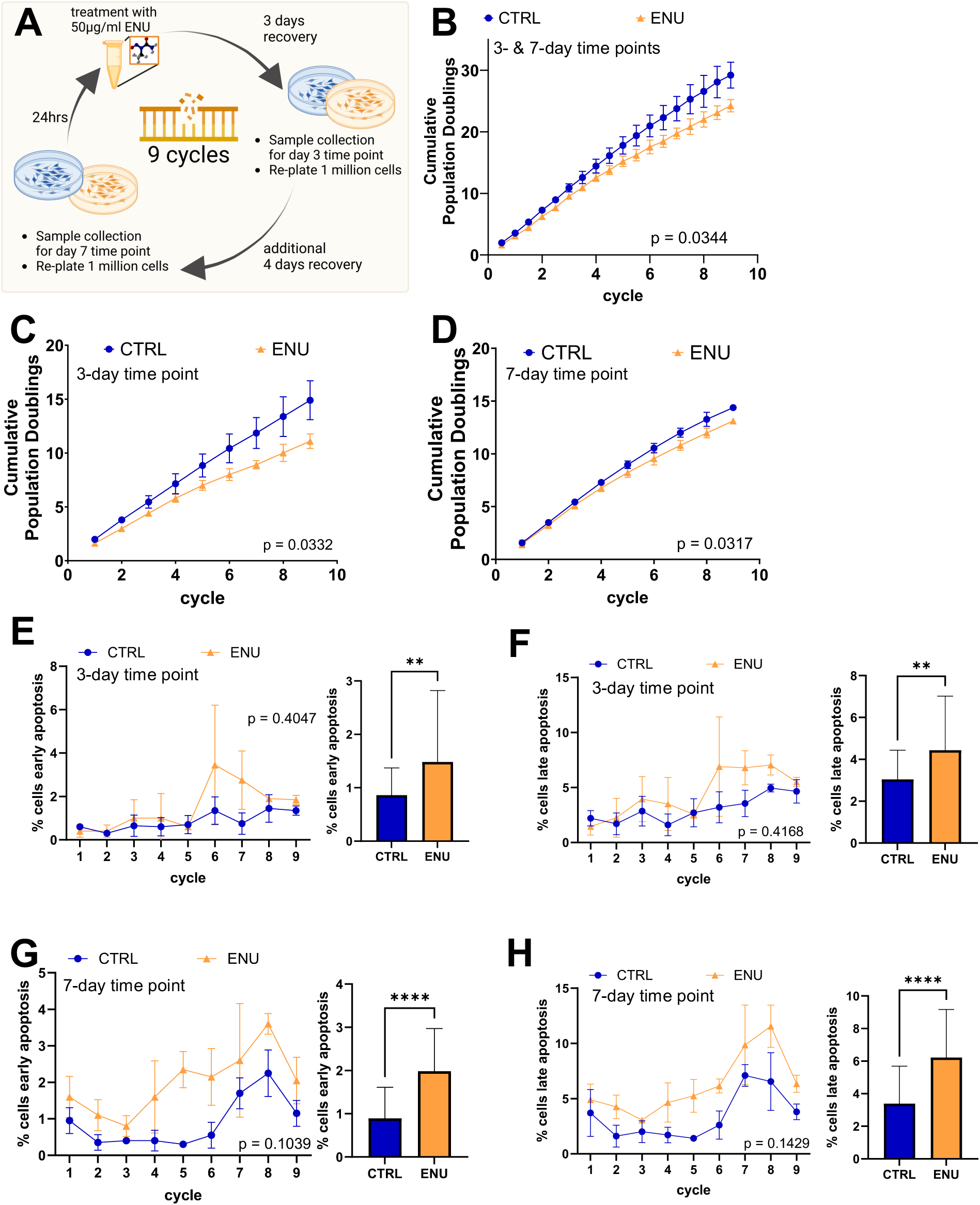
Repeated Mutagen Treatment of Normal Cells Only Slightly Affects Cell Growth and Death. **(A)** Schematic depiction of the experimental design. Human fetal lung fibroblasts (IMR-90) were treated with a sublethal dose of 50 µg/mL of N-ethyl-N-nitrosourea (ENU), which was followed by a recovery period of 7 days. At 3-day and 7-day time points, 1 million cells were passaged and a sample was taken for further analysis. This process was repeated for 9 cycles (Methods). **(B)** Cumulative population doublings (PDs) of the control (CTRL) and ENU-treated (ENU) groups throughout the experiment. Statistics: n=3; Data represent mean ± S.D.; P-value estimated using a two-way repeated measures ANOVA. **(C)** Cumulative PDs of the CTRL and ENU groups assessed at the 3-day time point. Statistics: Same as in (B). **(D)** Same as in (C), but at 7-day time point. Statistics: Same as in (B). **(E)** Percentage of cells detected in early apoptosis at the 3-day time point at each cycle (left) or with all cycles pooled (right). Statistics: n=2; Data represent mean ± S.D.; P-value of data from each cycle was estimated using a two-way repeated measures ANOVA. P-value of data from all cycles pooled was estimated using a paired t-test; **: p <= 0.005; ****: p<= 0.0001. **(F)** Same as in (E), but for late apoptosis. Statistics: Same as in (E). **(G)** Same as in (E), but for 7-day time point. Statistics: Same as in (E). **(H)** Same as in F, but for 7-day time point. Statistics: Same as in (E).

To analyze possible effects of a DNA damage response induced by ENU treatment in more detail, we performed bulk mRNA-sequencing analysis on samples collected at cycles 1 and 9 for the control group and cycle 9 for the ENU-treated group (Figures S1A-K; Table S1; Methods). As a positive control for the effect of cell passsaging, we first compared the control cycle 9 and the control cycle 1 groups, which resulted in 979 significantly differentially expressed genes (DEGs) (Figures 2A; Table S2)(*20*). We then performed targeted gene set enrichment analysis (GSEA) which showed that cell cycle-related genes were significantly decreased, while apoptosis-related genes were not significantly changed (Figures 2B-C). As cells are known to become senescent during passaging, we also tested for an enrichment of senescence-related genes using the SenMayo gene set(*21*), which were found to be significantly increased, as expected (Figure 2D). Furthermore, we found the DEGs to be positively correlated with DEGs found in a previous study on replicative senescence (Figure S1N).

**Figure 2.**
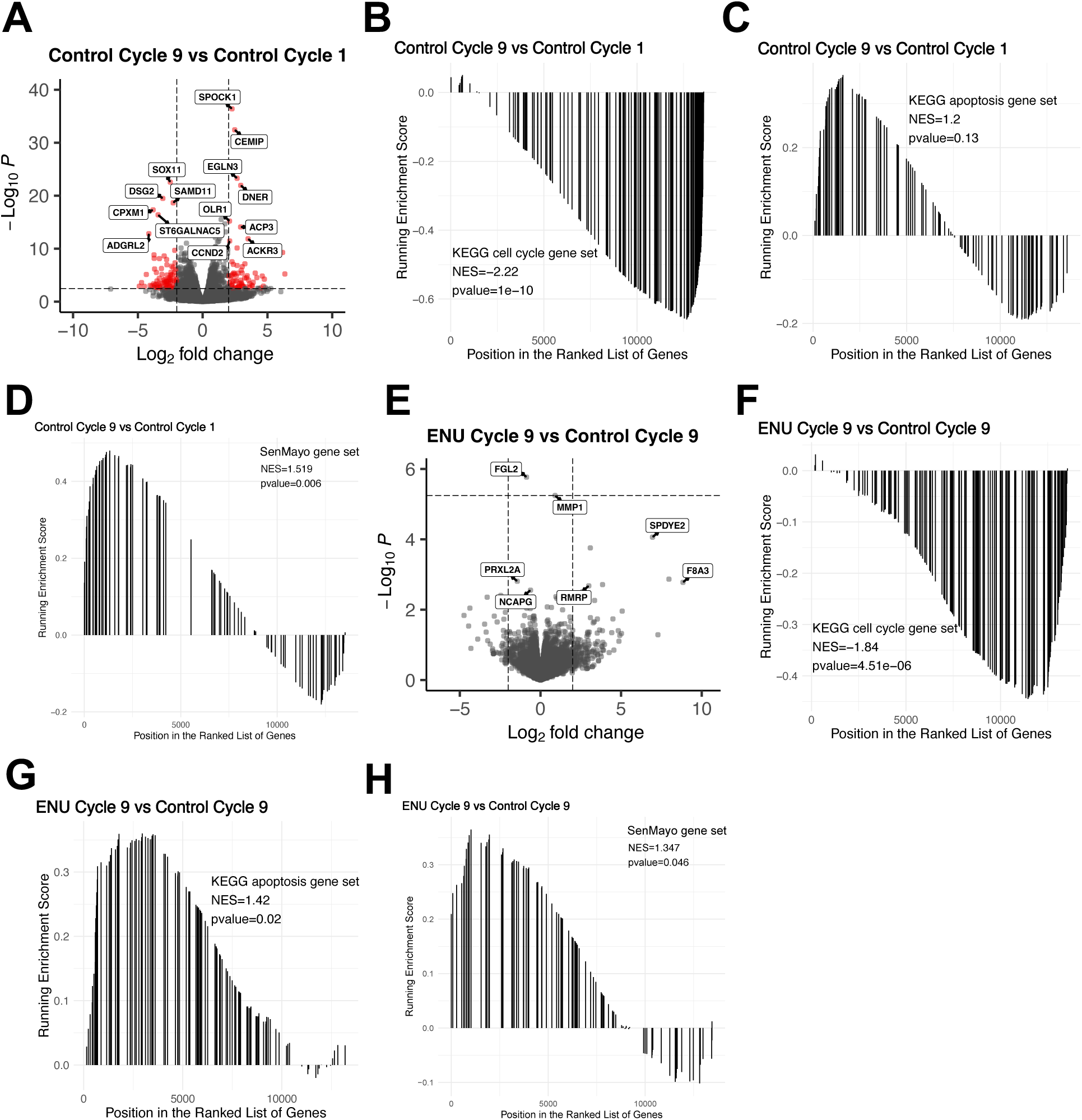
RNA-Sequencing Reveals Increased Cell Cycle Inhibition and Apoptosis in ENU-treated Cells. **(A)** Differential gene expression (DGE) of control cycle 9 vs. control cycle 1 groups. Red points indicate Log_2_ fold change>2 and adjusted p-value<0.05. See Table S1 for sample details and Table S2 for DGE results. Statistics: n=3; DGE p-values estimated using a Wald test followed by Benjamini-Hochberg correction for multiple hypothesis testing. **(B-D)** Gene set enrichment analysis (GSEA) of DGE results from the control cycle 9 vs. control cycle 1 comparison for KEGG cell cycle (B), KEGG apoptosis (C), and SenMayo senescence (D) gene sets. NES=normalized enrichment score. Statistics: n=3; p-values estimated using a permutation test. **(E)** Same as in (A), but for the ENU-treated cycle 9 vs. control cycle 9 comparison. See Supplementary Note 1 for batch correction note. Statistics: Same as in (A). **(F-H)** Same as in (B-D), but for the ENU-treated cycle 9 vs. control cycle 9 comparison. Statistics: Same as in (B-D).

To assess the effects of the repeated ENU treatment, we compared the ENU-treated cycle 9 and the control cycle 9 groups. Unexpectedly, this resulted in only 2 significant DEGs (Figure 2E; Table S2; Supplementary Note 1). However, after performing GSEA (which does not rely on p-value thresholds), we found that repeated ENU treatment had a similar, albeit weaker, effect as cell passaging – a significant decrease of cell cycle-related genes and a significant increase in senescence-related genes (Figure 2F, 2H). In contrast to the effect of cell passaging, we found a significant increase in the abundance of apoptosis-related genes in the ENU-treated cells relative to the controls (Figure 2G), confirming the results of the direct apoptosis measurements (Figures 1G-H).

Taken together, these phenotypic and molecular data suggest that the observed slight decrease in cell growth rate after repeated ENU treatment is, at least in part, due to DNA damage-inducd cell cycle inhibition and increased apoptosis. As the resulting effect sizes due to repeated ENU treatment (i.e. mutation accumulation) were relatively small, these results demonstrate the tolerence of normal primary cells proliferating in culture to this mutagen, which was previously known to have low cell toxicity when used at low doses(*22*).

### Repeated Mutagen Treatment of Normal Cells Results in Extremely High Mutation Burden

Somatic DNA mutations in normal (non-cancerous) cells, which are mostly unique to each cell, cannot be distinguished from sequencing artifacts after bulk whole genome sequencing. Hence, we used our established and highly accurate single-cell whole genome sequencing assay, which has been extensively validated by parallel analysis of kindred single-cell clones(*23, 24*), to analyze mutation burden in 21 single-cells from control and ENU-treated groups collected in 3 independent batches at cycles 3, 6, and 9 from the repeated ENU treatment experiment (Figure 1A, Table S3, Methods). Cells for mutation analysis were collected at the 7-day time point after each treatment cycle, as DNA damage responses should be abated by that time and all mutations fixed (Figure 2D). We then first corrected the mutations that were actually observed for the less than 100% genome coverage and sensitivity, which provided estimated mutation burdens higher than the observed burdens (Figures S2A-G; Methods). We also corrected for technical artifacts in the control group for cycle 1, which reduced mutation burden somewhat but had no effect at all on the ENU-induced mutation burden (Figures S3A-D; Methods). We found that the number of estimated SNVs per cell in the control group was initially 1,996±332 at cycle 1, which then increased to 19,064±823 following 3 ENU treatment cycles, (∼7.8-fold increase compared to cycle 1, p-value=1.54*10^-6^), and then 33,098±1,785 after 6 ENU treatment cycles (∼1.7-fold increase compared to cycle 3, p=1.90*10^-5^), and finally reached 55,954±4,066 after the 9th ENU treatment cycle (∼1.7-fold increase compared to the cycle 6, p =3.33*10^-9^) (Figures 3A and S2H). T>A and T>C substitutions, known to be induced by ENU(*22*), accounted for a large proportion of SNVs and showed the same trends as when analyzing the total mutation burden (Figures S2K-L). Importantly, there was no significant difference between control cells at cycle 1 and 9, ruling out that the increase in SNVs in ENU-treated cells was due to passaging. We also checked for the possibility of clonal expansion by looking at shared mutations between cells within each group, but only detected 8 SNVs that were shared between 2 cells in the ENU-treated group at cycle 6 (Figure S2F). This indicates a shared common ancestor, but not clonal expansion. This was expected as clones should not have the opportunity to grow over the relatively short number of cell divisions and the small number of individual cells analyzed. We conclude that our treatment paradigm increased mutational load to supraphysiological levels without serious toxicity, with the increase in SNVs over the course of the experiment occurring linearly in a cycle-dependent manner (Figure 3B).

**Figure 3.**
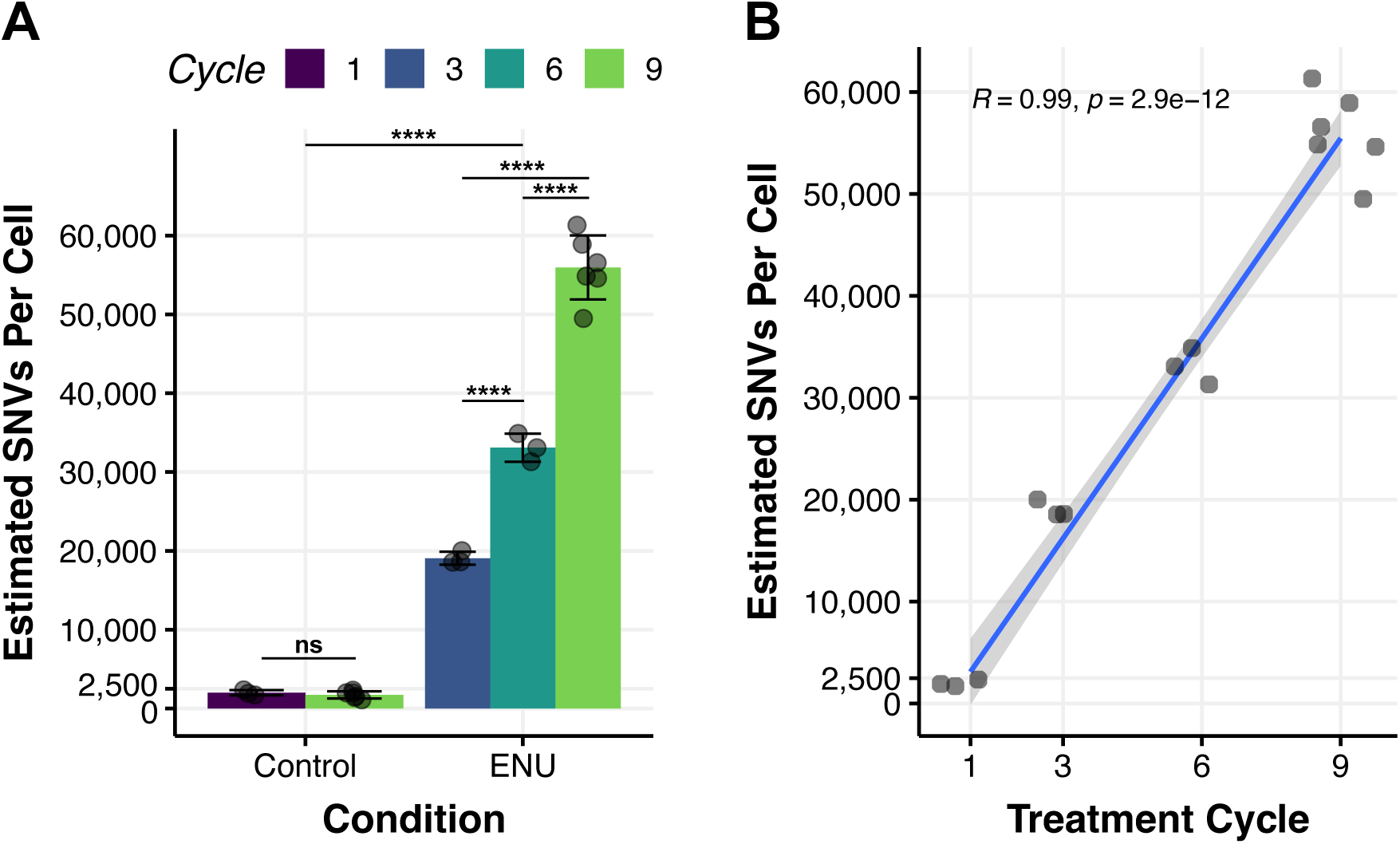
Repeated Mutagen Treatment of Normal Cells Results in Extremely High Mutation Burden. **(A)** Estimated load of somatic single nucleotide variants (SNVs) per cell after correction for genome coverage, sensitivity, and technical artifacts (Methods). See Figure S2H for observed SNV load per cell and Figures S2K-L for ENU-specific mutation burden. See Table S3 for sample details and Table S10 for mutation catalogue. Statistics: n=3 cells for cycles 1, 3, and 6; n=6 cells for cycle 9; data represent mean ± S.D. p-values of comparisons between cycles within the same condition were estimated with a two-sided linear model with estimated marginal means, Tukey’s HSD. P-values of comparisons of control and ENU-treated groups were estimated using a two-way ANOVA and shown at very top of plot. p-value legend: ns:p > 0.05, ****: p <= 0.0001. **(B)** Fit of linear model to estimated SNV load per cell shown in (A) vs. the treatment cycle cells were collected at. Note that cells from control cycle 1 are included and control cycle 9 is omitted. Shown at top is the fit of a linear equation. Statistics: data represent mean ± S.D.; Pearson R correlation coefficient and p-value of the correlation was estimated using a T-test.

While ENU is known to mainly induce base substitution mutations through alkylation damage, the estimated number of small insertions and deletions (INDELs) per cell was also observed to increase due to ENU treatment, starting from 123±49 in the control group at cycle 1, to 246±49 in the ENU-treated group at cycle 3 (∼1.8-fold increase, p=4.10*10^-3^); however, there was no further increase at cycles 6 and 9 (Figures S2I-J). As with SNVs, there was no significant difference in the amount of estimated INDELs between control cells from cycle 1 and cycle 9. INDELs have been previously observed to be induced by ENU, albeit at low frequencies(*25*), and could be caused by DNA polymerase slippage during DNA replication at sites of alkylation damage. While the lack of an increase in INDELs after cycle 3 in the ENU-treated group may be suggestive of negative selection acting to prevent a further increase, as was seen in a recent study (*26*), our current finding remains inconclusive due to the low amount of observed INDELs per cell (∼20 INDELs per cell).

We then characterized the underlying SNV mutational processes in the aggregated mutations from the cells at all cycles within the control and ENU-treated groups (Methods). The ENU-treated group showed an expected increase in the relative contribution of thymine-to-adenine (T>A), followed by thymine-to-cytosine (T>C) base transversions (Figures 4A, S2K, S3A). This is in keeping with previously published ENU-induced mutational spectra by us and others(*22, 27*). We then extracted SNV mutational signatures *de novo* using non-negative matrix factorization with a reference set of mutations from previous single-cell whole genome studies (*23, 28–30*) (Figures S3E-F) and identified two signatures, labeled as SBSA and SBSB, which were dominant in the cells of the ENU-treated and control groups, respectively (Figures 4B and S3G). Importantly, we found that SBSA resembled the SIGNAL human signature associated with ENU-treatment, validating the source of the induced mutations (Figure 4C)(*31*). We then compared our *de novo* signatures to those in the COSMIC database and found that SBSB was most similar to SBS5 a ‘clock-like’ mutational signature that has been previously found to correlate with cellular replication during aging (*28, 32*) (Figure 4D). Though relatively few INDELs were observed (369 in total), we also provide a characterization of their mutational spectra (Figure S3H; Supplementary Note 2).

**Figure 4.**
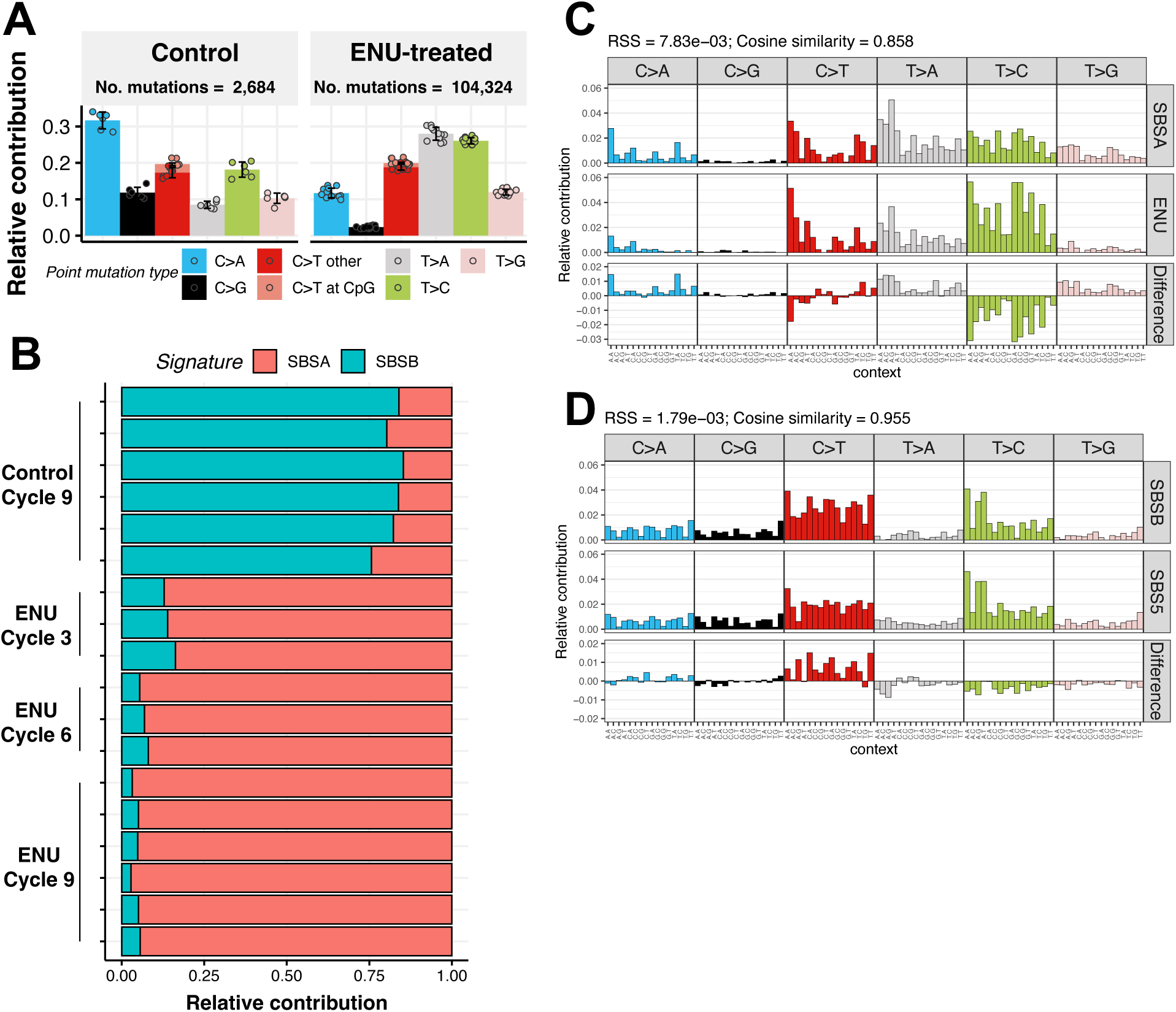
Mutational Spectra And Signatures Resulting From Repeated Mutagen Treatment. **(A)** Mutational spectra of the relative contribution of SNV types per cell grouped by condition. See Figure S3A for groups stratified by cycle. Note that cells from the control cycle 1 group were excluded from the mutational spectra and signatures analyses due to abnormally high C>T frequency suggestive of a sample preparation artifact (Figures S3A-D; Methods). Statistics: n=3 cells for cycles 3 and 6; n=6 cells for cycle 9; Data represent mean ± S.D. (**B)** Relative contribution of de novo mutational signatures SBSA and SBSB to individual cells (shown as rows) grouped by condition and cycle. See Figure S3F for determination of number of signatures to extract and Figure S3G for absolute contribution. **(C)** Comparison of de novo mutational signature SBSA to the ENU signature from the SIGNAL database. Statistics: cosine similarity 0.858, residual sum of squares=7.83*10^-3^. **(D)** Comparison of de novo mutational signature SBSB to most similar fitted signature SBS5 from the COSMIC database. Statistics: cosine similarity=0.955, residual sum of squares= 1.79*10^-3^.

### Negative Selection Against ENU-Induced Accumulation of Damaging Coding and Non-Coding Variants

The above results show that the mutation burden of primary cells after 9 cycles of ENU treatment increased to an estimated ∼56,000 SNVs per cell (i.e., ∼10 times the SNV burden typically found in normal cells from aged tissues(*6*)), with only a slight effect on cell growth and death. From this, we hypothesized that the sustained cell survival and growth in the face of a very high mutation burden could be explained by negative selection against harmful mutations. Our dataset presented a unique opportunity to test this hypothesis as there were a total of 108,600 SNVs observed collectively in all cells. Note that this was smaller than the estimated number after corrections for sensitivity and genome coverage as mentioned above (Methods), but still expected to provide sufficient power for statistical testing of selection pressure (Figure S2H).

To obtain a random background distribution of expected mutations against which to compare the observed mutations in testing for selection pressure, we used the SigProfilerSimulator tool (*33*). This method controls for variation in genomic coverage, mutational signatures, and sequence context of the observed mutations. With this, we simulated 10,000 instances of random mutations for each treatment cycle in both the control and ENU conditions (Methods). Following this, both observed and expected SNVs in coding regions were annotated as synonymous (non-protein-altering) or non-synonymous (protein-altering) variants, using the Ensemble Variant Effect Predictor program (Table S4)(*34*). The simulations of expected mutations performs well in predicting the number of synonymous variants (R=0.94, p=3.6*10^-10^; Figure S4A), which should be under minimal selection. We then measured selection pressure within each group by calculating the Log_2_(observed/expected) (written as Log_2_(O/E) hereafter) ratios for each variant type using the frequency of observed and expected variants (*35*). The direction of selection pressure (i.e. neutral, positive, or negative) was ascertained by comparing the resulting Log_2_(O/E) ratio to 0, where no difference from 0 is what would be expected under neutral selection (*36*) (Methods). We then used bootstrapping to determine the threshold for the frequency of observed variants needed to obtain sufficient power for statistically significant results (p<0.05) at various Log_2_(O/E) ratios (Figures S4B; Table S5; Methods). Finally, to account for the known effect of DNA repair in depleting the frequency of mutations within transcribed regions through transcription-coupled repair (TCR) and mismatch repair (MMR) (*37, 38*), we normalized the Log_2_(O/E) ratios of variants within transcribed regions by the Log_2_(O/E) ratio of mutations within intronic regions (assumed to be equally transcribed, but under neutral selection; Methods).

With this approach, we found that while there were significant cycle-dependent increases in both synonymous and non-synonymous variant frequencies in the ENU-treated group relative to the controls (Figure 5A), the Log_2_(O/E) ratios of non-synonymous variants at cycles 6 and 9 in the ENU-treated group indicated that the frequency of these damaging variants was significantly lower than what would be expected, indicating negative selection (Figures 5B and S4C-E, right panels). The negative Log_2_(O/E) ratios of non-synonymous variants at cycles 6 and 9 corresponded to a ∼20% reduction in mutation burden, equivalent to a decrease of 14.5 ± 9.1 and 19.9 ± 12.5 variants per cell at cycles 6 and 9, respectively (Figure 5C). In regards to synonymous variants, the Log_2_(O/E) ratios of these variants trended towards 0 across all cycles in the ENU-treated group, which would suggest neutral selection (as would be expected for these benign variants). However, the frequency of observed synonymous variants was below the threshold set by the previously described power analysis, which prevented us from making any definite conclusions (Figures 5B and S4C-E, left panels; Table S5).

**Figure 5.**
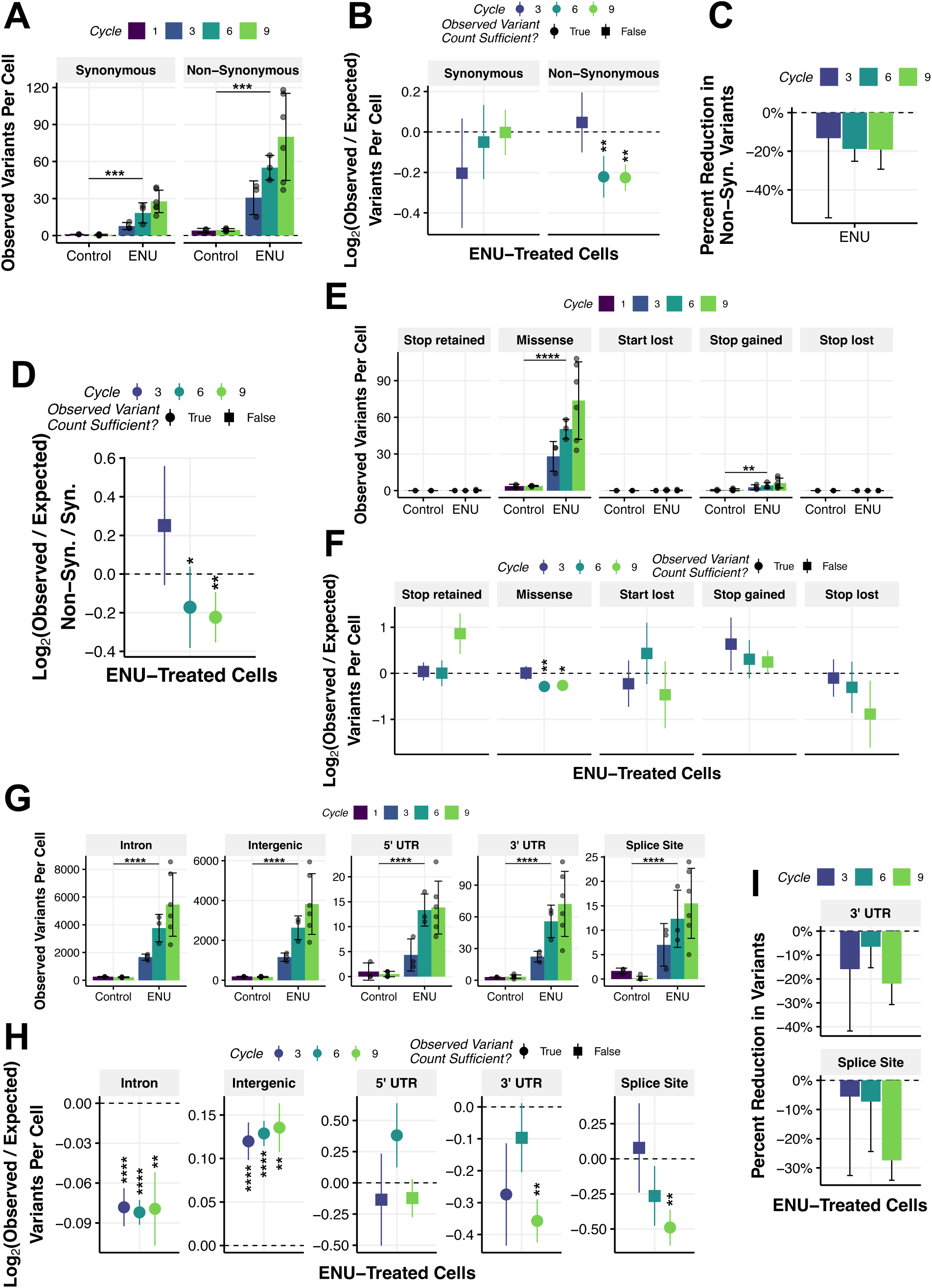
Negative Selection Against Damaging Coding and Non-Coding Variants in ENU-treated Cells. **(A)** Observed frequency of synonymous and non-synonymous variants per cell. See Table S3 for sample details and Table S5 for variant annotation. Statistics: n=3 cells for cycles 1, 3, and 6; n=6 cells for cycle 9; data represent mean ± S.D; P-values of comparisons of control and ENU-treated groups were estimated using a two-way ANOVA and shown at very top of plot; p-value legend: ***: p <= 0.001. **(B)** Log_2_(O/E) ratio of synonymous and non-synonymous variants for ENU-treated cells. See figures S4C, S4D, and S4E for results of without normalization, results of control cells, and results per cell, respectively. See Table S5 for detailed results. Statistics: n=3 cells for cycles 3 and 6; n=6 cells for cycle 9; data represent mean ± S.D; P-values to assess if the Log_2_(O/E) ratio significantly deviates from 0 were estimated using a permutation test (n=10,000) but were only estimated if the frequency of observed variants allowed for sufficiency statistical power (Figure S4B; Table S5); p-value legend: **: p <= 0.01. **(C)** Estimated percent of non-synonymous variants out of total coding variants that are reduced in the ENU-treated group due to negative selection. Statistics: n=3 cells for cycles 3 and 6; n=6 cells for cycle 9; data represent mean ± S.D. **(D)** Log2O/E of the N/S ratio for ENU-treated cells. See figures S4I and S4J for results of control cells and per cell, respectively. Statistics: Same as in (B). p-value legend: *: p <= 0.05; **: p <= 0.01. **(E)** Observed frequency of various types of non-synonymous variants per cell. Statistics: Same as in (A). p-value legend: ****: p <= 0.0001. **(F)** Log2O/E of various types of non-synonymous variants for ENU-treated cells. See figures S4F, S4G, and S4H for results of without normalization, results of control cells, and results per cell, respectively. Statistics: Same as in (B). p-value legend: *: p <= 0.05; **: p <= 0.01. **(G)** Observed frequency of various types of non-coding variants per cell. Statistics: Same as in (A). ****: p <= 0.0001. **(H)** Log_2_O/E of various types of non-coding variants for ENU-treated cells. See figures S5K, S5L, and S5M for results of control cells, without normalization, and per cell, respectively. See Supplementary Note 5 for discussion on results of the intergenic region. Statistics: Same as in (B). ****: p <= 0.0001; **: p <= 0.01. **(I)** Estimated percent reduction of variants in 3’ UTR and splice region variants that are protected against in the ENU-treated group due to negative selection. Statistics: Same as in (C).

We then sought to verify our finding of negative selection against non-synonymous variants by using a method which calculates dN/dS ratios (i.e. the dNdScv program), an alternative measure of selection commonly used in cancer or evolutionary studies (Methods)(*39*). This metric compares the observed number of non-synonymous mutations (missense, nonsense, and splice-site) to the number expected under neutral evolution, using synonymous mutations as a neutral reference. A global dN/dS ratio significantly greater than 1 indicates positive selection, while a ratio less than 1 suggests negative selection. As this is effectively the same as calculating the O/E of non-synonymous mutations over the O/E of synonymous mutations, we calculated the Log_2_(O/E) non-synonymous/synonymous (N/S) to facilitate the comparison. Synonymous variants, assumed to be functionally neutral, occur in the same sequence contexts and at the same depth as non-synonymous variants, so they serve as an internal baseline for the true mutation rate introduced by ENU. A deficit of non-synonymous variants relative to this neutral baseline is therefore attributable to negative selection rather than to sequencing bias or differences in mutagenesis. Log_2_(O/E) (N/S) ratios were significantly below 0 at cycles 6 and 9 in the ENU-treated group (Figures 5D, S4F-G), indicating negative selection and corroborating our previous result (Figure 5B). Before reviewing the results of the dNdScv method, it should be noted that there are important differences between this method and our O/E method. Unlike our O/E method, which empirically estimates expected mutations via genome-wide simulations accounting for coverage and mutational signatures (*35*), the dNdScv method relies on probabilistic modeling to estimate the expected mutations, assumes uniform genomic coverage, and expects abundant mutations in coding regions. Therefore, the assumptions of this method make it less suitable for our single-cell whole genome sequencing data type, i.e., variable genome coverage and a limited number of coding mutations (Figures S2A and 5A). Expectedly, global dN/dS ratios were in contrast to Log_2_(O/E) N/S ratios (Figure S5A; See Supplementary Note 3 for a detailed comparison between methods).

To identify the contribution of the various types of non-synonymous variants to the negative selection signal observed in the ENU-treated group, we stratified the non-synonymous variants into stop retained, missense, start lost, stop gained, and stop lost variants and calculated their respective Log_2_(O/E) ratios (See Supplementary Note 4 for details on stop lost variants). This revealed that missense variants made up the majority of non-synonymous variants in the ENU-treated group (Figure 5E) and also that the negative selection signal was driven almost exclusively by these variants (Figures 5F and S4H-J).

Next, we tested if variants in non-coding regions were under selection pressure. For this, we again used the Variant Effect Predictor program to annotate SNVs in non-coding regions, i.e., introns, intergenic, 5’ untranslated regions (UTR), 3’ UTR, and splice sites, and then quantified their frequencies (Table S4, Methods)(*34*). As with the coding variants, non-coding variants showed cycle-dependent increases in the ENU-treated group relative to the controls, with variants in intronic and intergenic regions accounting for 49.9% and 35.1% of all non-coding variants, respectively (Figure 5G). For ENU-treated groups at all cycles, intronic region variants Log_2_(O/E) ratios ≈ −0.08, significantly less than what would be expected by chance (Figure 5H, Table S5). Since the vast majority of introns (with the exception of splice sites and other conserved non-coding elements) do not contain functional sequences and are evolving at neutrality in the germline, but are still subject to DNA repair such as TCR and MMR(*37, 38*), we interpret the negative Log_2_(O/E) ratios in these regions as a signal of DNA repair rather than negative selection. In fact, as described at the beginning of this section, we use the intron Log_2_(O/E) ratio to account for the confounding effect of DNA repair on the Log_2_(O/E) ratio. In intergenic regions for ENU-treated groups at all cycles, we found that the Log_2_(O/E) was ∼0.13, significantly greater than what would be expected by chance and indicative of mutation enrichment (Figure 5H, Table S5). Similar to introns, the vast majority of intergenic regions lack functional sequences and are evolving at neutrality in the germline. However, DNA repair activity in these regions is known to be substantially reduced and likely explains the positive Log_2_(O/E) ratios(*40, 41*). See also Supplementary Note 5.

Out of all the non-coding variants, only Log_2_(O/E) ratios of variants occurring in 3’ UTR and splice regions in the ENU-treated group at cycle 9 were significantly less than 0 (Figures 5H, S4K-M, Table S5), which was indicative of negative selection and supported by previous studies on germline selection pressures(*40, 42, 43*). The Log_2_(O/E) ratios of 3’ UTRs and splice sites in the ENU-treated group at cycle 9 corresponded to ∼20-25% reduction in mutation burden, a decrease of 19.3±12.5 and 6.2±2.7 variants per cell, respectively (Figure 5I).

Taken together, these results show that potentially damaging variants in both coding and a subset of non-coding regions did not accumulate to levels that would be expected in the context of the extremely high mutation burden induced by ENU treatment, and instead were subject to negative selection, resulting in lower mutation burdens within critical genes supporting essential cellular functions *in vitro*. This would explain how the population of cells exposed to repeated mutagen treatments was able to avoid a dramatic decline in growth rate, even as overall SNV burden continued to increase linearly throughout the experiment. Of note, the selection pressure analysis was limited to mutations that were actually physically detected, which are ∼20% of the ∼56,000 estimated mutations at cycle 9 in the ENU-treated group (Figures 3A and S2H). Hence, these results have relevance for the much smaller numbers of SNVs that are estimated to accumulate during normal aging, e.g., from ∼3,000 in human B lymphocytes to ∼ 9,000 in human hepatocytes(*6*).

### Pathways Supporting Fibroblast Growth and Cell Identity are Protected from Mutation Accumulation

To gain insight into selective pressures at broader levels associated with cellular function, we investigated if gene sets active in proliferating human fibroblasts were under negative selection. To do this, we analyzed the SNVs across annotated functional genomic regions relevant to human lung fibroblasts (i.e. exons, upstream promoters, and enhancers) that were grouped into various gene sets(*44–46*). We then calculated the frequency of SNVs within these regions as well as the Log_2_(O/E) ratios in a similar manner to the previous analyses (Methods).

First, we analyzed non-expressed genes and expressed genes as determined by our bulk mRNA-sequencing data (Figures S1A-B; Table S7; Methods). In keeping with the previous analyses of other regions (e.g. exonic and intronic regions), we found cycle-dependent increases of SNV frequency in non-expressed and expressed regions for the ENU-treated group relative to the controls (Figure 6A). Log_2_(O/E) ratios of SNVs in expressed genes were significantly below 0 at all cycles in the ENU-treated group, indicative of negative selection (Figures 6B and S6A-C, right panels). In contrast, the Log_2_(O/E) ratios of SNVs in non-expressed genes did not significantly deviate from the expectation in the ENU-treated group at cycle 9, indicating neutral selection (Figures 6B and S6A-C, left panels). Next, we further subdivided the expressed genes into nonessential and essential gene sets based on previous *in vitro* screens (Table S7) and repeated the same analysis as above (*47, 48*). This showed increased SNV frequency in the ENU-treated group relative to the controls (Figure 6C), along with a Log_2_(O/E) ratio significantly below 0 at cycle 9 in the ENU-treated group, indicative of negative selection (Figures 6D and S6D-F, right panels). Of note, this is in agreement with global dN/dS ratios on truncating mutations across 17 genes essential from a recent large-scale study sampling the normal oral epithelium(*49*).

**Figure 6.**
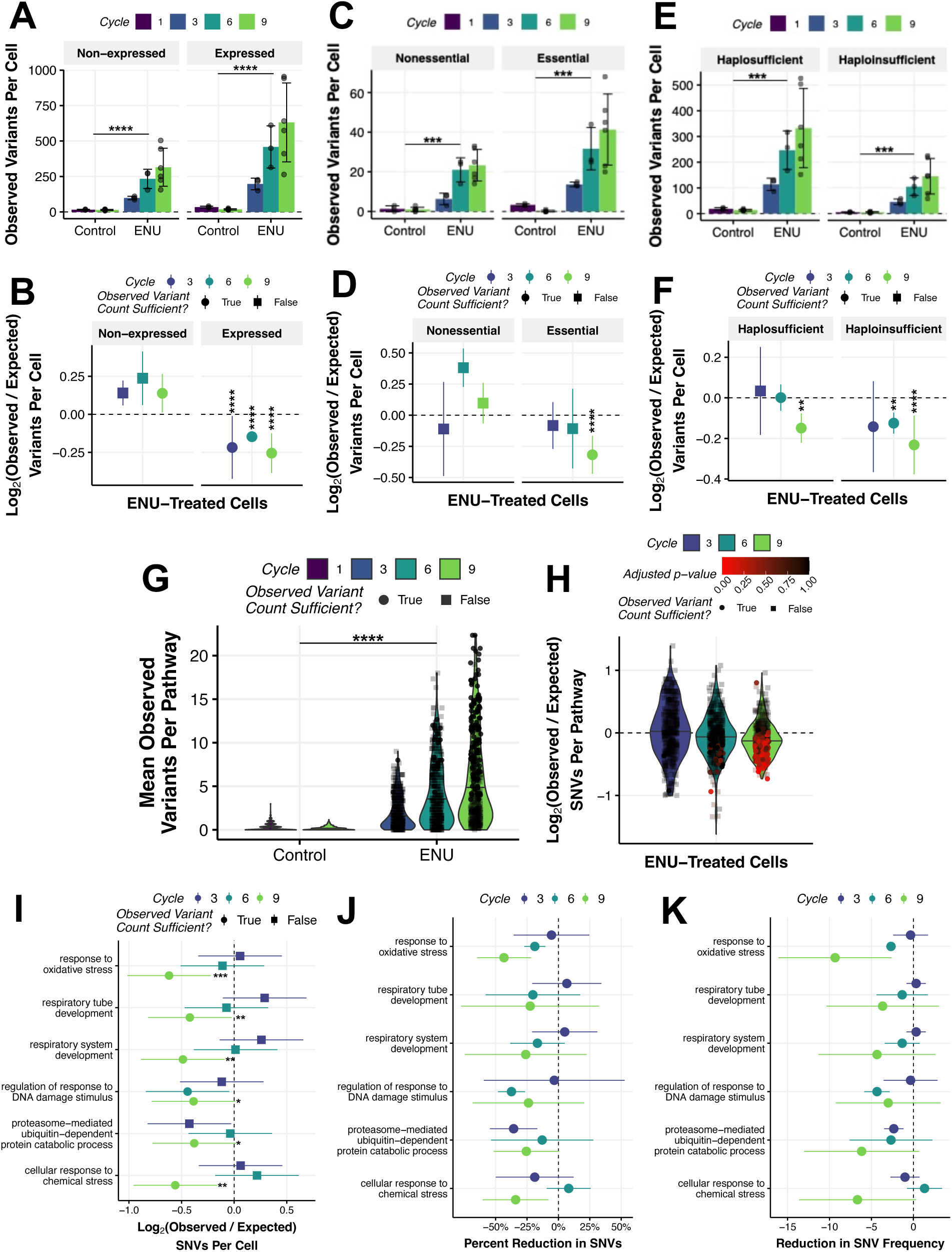
Pathways Supporting Fibroblast Growth and Cell Identity are Protected from Mutation Accumulation. **(A)** Observed frequency of SNVs per cell in functional regions associated with genes that were determined to be non-expressed or expressed from bulk mRNA transcriptome data (Table S7). Statistics: n=3 cells for cycles 1, 3, and 6; n=6 cells for cycle 9; data represent mean ± S.D; P-values of comparisons of control and ENU-treated groups were estimated using a two-way ANOVA; p-value legend: ****: p <= 0.0001. **(B)** Log_2_(O/E) ratio of SNVs in functional regions associated with genes determined to be non-expressed or expressed expressed in bulk mRNA transcriptome data (Table S7). See figures S6A, S6B, and S6C for results without normalization, results of control cells, and results per cell, respectively. Statistics: n=3 cells for cycles 3 and 6; n=6 cells for cycle 9; data represent mean ± S.D; P-values to assess if the Log_2_(O/E) ratio significantly deviates from 0 were estimated using a permutation test (n=10,000) but were only estimated if the frequency of observed variants allowed for sufficiency statistical power (Figure S4B; Table S5); p-value legend: ****: p <= 0.0001. **(C)** Observed frequency of SNVs per cell in functional regions associated with nonessential or essential gene sets expressed in bulk mRNA transcriptome data (Table S7). Statistics: Same in in (A); p-value legend: ***: p <= 0.001. **(D)** Log_2_(O/E) ratio of SNVs in functional regions associated with nonessential or essential gene sets expressed in bulk mRNA transcriptome data (Table S7). See figures S6D, S7E, and S7F for results without normalization, results of control cells, and results per cell, respectively. Statistics: Same as in (B); p-value legend: ****: p <= 0.0001. **(E)** Observed frequency of SNVs per cell in functional regions associated with haplosufficient or haploinsufficient gene sets expressed in bulk mRNA transcriptome data (Table S7). Statistics: Same in in (A); p-value legend: ***: p <= 0.001. **(F)** Log_2_(O/E) ratio of SNVs in functional regions associated with haplosufficient or haploinsufficient gene sets expressed in bulk mRNA transcriptome data (Table S7). See figures S6G, S6H, and S6I results without normalization, results of control cells, and results per cell, respectively. Statistics: Same as in (B); p-value legend: **: p <= 0.01; ****: p <= 0.0001. **(G)** Mean observed frequency of SNVs within the functional regions of pathways expressed in bulk mRNA transcriptome data (Table S8). See figure S6J for top 20 most enriched pathways. Statistics: n=399 pathways; P-values of comparisons of control and ENU-treated groups were estimated using a two-way ANOVA; p-value legend: ****: p <= 0.0001. **(H)** Mean Log_2_(O/E) ratio of SNVs within the functional regions of pathways expressed in bulk mRNA transcriptome data (Table S8). See figure S6K and S6L for results of control cell and results without normalization, respectively. See table S9 for detailed results. Statistics: n=399 pathways; Otherwise, same as in (B), with the addition of Benjamini-Hochberg correction for multiple hypothesis testing of pathways. **(I-K)** Mean Log_2_(O/E) ratio (I), estimated percent reduction in SNVs (J), and estimated reduction in SNV frequency (K) within functional regions of pathways found to be under significant negative selection (p <= 0.05) at cycle 9 in the ENU-treated group. Statistics: Same as in (B); ; p-value legend: *: p <= 0.05; **: p <= 0.01; ***: p <= 0.001.

As negative selection is expected to be generally weak due to the diploid nature of the human genome(*39, 50*), but stronger in haploinsufficient regions (i.e. regions where both alleles are required for function), we subdivided expressed genes into haplosufficient (pLI > 0.1) and haploinsufficient (Table S7) (pLI < 0.9) gene sets based on a large scale study of protein-coding genetic variation(*35*). As expected, analysis of these gene sets showed increased SNV frequency in the ENU-treated group relative to the controls (Figure 6E). Log_2_(O/E) ratios were significantly below 0 in both haplosufficient and haploinsufficient gene sets for the ENU-treated cells at cycle 9. However, only the ENU-treated group at cycle 6 showed Log_2_(O/E) ratios significantly below 0 for the haploinsufficient gene set and not the haplosufficient gene set, suggesting overall stronger negative selection of haploinsufficient genes, as would be expected (Figures 6F and S6G-I). In further support of this, when the Log_2_(O/E) was compared between haplosufficient and haploinsufficient gene sets for ENU-treated cells at cycle 9, we found that the Log_2_(O/E) ratio was considerably lower for the haploinsufficient genes (−0.37±0.16) as compared to the haplosufficient genes (−0.28±0.04). In summary, these results suggest that damaging SNVs are selectively purged from functional genomic regions based on expression status, essentiality, and dosage sensitivity, extending our previous results on negative selection of damaging SNVs in coding and non-coding regions.

Moving to the pathway level, we identified 399 gene ontology (GO) biological processes as significantly enriched in the set of expressed genes from our bulk RNA-seq data (Figure S6J, Table S8, Methods). As expected, the mean frequency of SNVs within each pathway showed significant cycle-dependent increases of SNV frequency in the ENU-treated group relative to the controls (Figure 6G). At the global level, Log_2_(O/E) ratios of the identified pathways in the ENU-treated group showed a cycle-dependent decrease, suggesting widespread negative selection at the pathway level (Figures 6H, S6K-L, Table S9). At cycle 9 in the ENU-treated group, 6 pathways showed Log_2_(O/E) ratios significantly below 0 after multiple hypothesis correction, indicative of negative selection. The pathways under negative selection were characterized as those related to basic cellular processes supporting lung fibroblast cell growth *in vitro* and cellular identity, i.e., ‘response to oxidative stress’ (GO:0006979), ‘respiratory tube development’ (GO:0030323), ‘respiratory system development’ (GO:0006979), ‘regulation of response to DNA damage stimulus’ (GO:2001020), ‘proteasome-mediated ubiquitin-dependent protein catabolic process’ (GO:0043161), and ‘cellular response to chemical stress’ (GO:0062197) (Figures 6I-K, Table S9). To ensure that the pathways we had identified to be under negative selection were not false positives, we repeated the pathway analysis, but employed bootstrapping to randomly shuffle genes amongst pathways before calculating the Log_2_(O/E) ratios (Figures S7A-C, Methods). The results of this served as a negative control, which had no significant overlap with the pathways we identified to be under significant negative slection pressure, providing a validation of our results (Figures S7D-E).

In summary, these results suggest that in the context of very high mutation burden, pathways supporting oxidative stress, cellular identity, and protein homeostasis in proliferating cells *in vitro*, are protected from the accumulation of SNVs by negative selection. Importantly, these pathways are biologically significant, as they appear to be relevant to the conditions of *in vitro* cell culture and to the unique charecteristics of lung fibroblasts, highlighting the functional importance of preventing mutation accumulation. These analyses show how negative selection allows a population of dividing cells to survive supraphysiological mutation burden and underscores the impact of random somatic SNVs on essential cellular functions.

## Discussion

Somatic mutations are known since the 1970s to be the main cause of cancer(*51*). Mainly because of the exponential increase in cancer risk with age(*52*), aging itself has been suggested to be caused by the accumulation of somatic mutations(*3, 8, 53*). Testing this hypothesis has long been constrained by a lack of methods to quantitatively analyze somatic mutations in normal tissues during aging. However, both single-cell and single-molecule methods have recently been developed to accurately quantify randomly accumulating SNVs and INDELs. Using these methods, we and others have demonstrated that somatic mutation burden is far higher than expected based on the known germline mutation frequency and increases with age up to several thousands of mutations per cell in aged subjects depending on the tissue(*6, 9*). However, a causal relationship of increased mutation burden and age-related functional decline and disease risk remains untested. Instead, attention has been focused on clonally amplified somatic mutations and their causal relationship to non-cancerous, age-related diseases(*15, 54*).

In the present study, we tested for a possible limit to somatic mutation accumulation in a cell culture model, in which high levels of somatic mutations were induced by multiple low doses of the powerful mutagen ENU (Figure 1A). In this system, we were able to linearly increase the somatic mutation burden to ∼56,000 SNVs per cell, over 30-fold higher than untreated control cells (Figure 3A). Remarkably, this supraphysiological mutation load only slightly impacted cell growth and death rates (Figures 1B and 1E-H), suggesting that most cells can sustain an astounding level of genetic information loss. The most straightforward explanation for these findings is that the mammalian genome is highly robust, and that even very high numbers of point mutations are unlikely to adversely affect cellular function. Indeed, this would be supported by results from a recent study in which as many as ∼30,000 SNVs were observed in intestinal crypts of elderly patients with genetic defects in DNA mismatch repair(*55*).

An alternative explanation for the high tolerance of actively proliferating cells to mutation accumulation is negative selection against mutations in genomic regions important for cell survival. Evidence for negative selection against somatic mutations has been found in tumors(*56–58*). However, in cancer, positive selection is the dominant force, arising from mutations in cancer driver genes(*39*). Evidence for negative selection of somatic mutations in normal cells is scarce because of the small numbers of mutations available as compared to tumors, which limits robust statistical testing(*6*). Our present analysis of untreated control cells indeed suffered from this same limitation, and we were unable to convincingly demonstrate negative selection of mutations in these cells (Figure S4D, Table S5). However, our ENU-treated cells did yield a sufficiently high number of SNVs to test for negative selection (Figure S2H). With this, we found multiple lines of evidence for negative selection acting on potentially deleterious coding and non-coding variants. While the frequency of both synonymous and non-synonymous variants increased linearly with ENU treatment (Figure 5A), deeper analysis using observed vs. expected (Log_2_(O/E)) ratios revealed a significantly lower frequency of high impact, non-synonymous variants, especially missense variants, in the ENU-treated cells as compared to our null model (Figures 5B-C). This suggests that an actively proliferating population of fibroblasts has the capacity to selectively purge the most harmful coding mutations from their genomes, preserving cellular fitness despite the extremely high number of random mutations accumulating genome wide, and agrees with the selection pressures found from analyzing germline mutations(*35, 59–61*). This main finding was recently validated in an *Msh2^-/-^* mouse model, which found negative selection against somatic stop lost variants(*26*). Additionaly, negative selection was also detected in non-coding regions for variants occurring in 3’ UTR and splice sites (which collectively made up ∼1.1% of all the observed non-coding variants) (Figures 5G-I), which is in agreement with recent literature analyzing selective pressures in these regions (*43*).

Our analysis of gene sets and pathways revealed negative selection to also be acting at a broader level, highlighting the biological significance. In the primary lung fibroblasts treated with ENU, we found evidence of negative selection in functional genomic regions of expressed, essential and haploinsufficient genes (Figures 6A-F). This finding was recently recapitulated in a population scale study on somatic mutations in oral epithelium cells that observed negative selection on essential genes, which overlaps with the sets of essential and haploinsufficient genes that was used in our study (Figures 6C-F)(*62*). Moreover, signals of negative slection were identified in actively expressed pathways, such as oxidative stress, respiratory system development, and response to DNA damage (Figures 6I-K), which were biologically relevant to our *in vitro* experimental conditions and the lung fibroblast cell type that was used. For example, selection was observed against genes involved in respiratory system development, which include genes such *FGF10, FOXF1,* and *COL3A1* that are known to be crucial for lung fibroblast identity and function(*63–65*). This suggests that despite the massive somatic mutational burden across the genome overall, random somatic mutations that occur in pathways critical for cellular survival do indeed have a damaging effect.

Our work provides conclusive evidence that very high numbers of somatic mutations, collectively, may have a functional impact on normal cells *independent* of clonally amplified mutations. However, because a supraphysiological amount of mutations was required to detect negative selection, this puts into question whether there is an effect of negative selection on the much lower numbers of somatic mutations that accumulate during aging. In this respect, it should be noted that the actual number of SNVs physically observed wass much smaller that the ∼56,000 estimated based on genome coverage and sensitivity, i.e., no more than ∼12,000 (Figures 3A and S2H). This is not that much above the ∼2,000 and ∼5,000 SNVs per cell, depending on the cell type analyzed, that have been estimated to have accumulated at old age(*6*). Moreover, we did not observe an association with the strength of negative selection and mutation burden (Figure 5B). Hence, it is conceivable that negative selection is a general mechanism to eliminate somatic mutations that adversely impact cell function, similar to purifying selection in the germline.

Another important aspect when considering the implications of negative selection are the differences between our *in vitro* model and the *in vivo* context. While negative selection in our model acted on cell proliferation, survival, and apoptosis, immune surveillance (i.e. natural killer cells and macrophages) would provide additional targets for *in vivo* elimination of fibroblasts displaying neoantigens or excreting senescence-associated factors. Moreover, in normal adult subjects, cell division rates are very low with few exceptions, most notably the lymphoid and intestinal systems(*19*). In proliferating cells, such as human B lymphocytes, we have previously demonstrated that SNVs accumulate with age much slower in the functional part of the genome than in the genome overall, indicating negative selection of mutations arising during the process of B cell aging(*66*). By contrast, somatic mutations accumulating in non-dividing adult cells, such as neurons or cardiomyocytes, cannot be selected against through proliferation or apoptosis, which is evidenced by excess accumulation of damaging mutations(*12, 67*). This suggest that age-related somatic mutation accumulation would have the greatest impact on non-proliferative cells, which may be a determinant in species lifespan as predicted by the disposable soma theory(*68*).

As a limitation of our study, we should point out that we only analyzed SNVs, and the few small insertions and deletions induced by ENU. However, SNVs are much less likely to have a functional impact than genome structural variants (SVs), including large deletions, translocations, and copy number variants. Indeed, this is clearly indicated for INDELs, which in our present study did not linearly accumulate during the ENU treatment (Figures S2I-J). In support of this, we recently conducted a study where the limit to the number of INDELs, ∼16,000 per cell, was determined by profiling somatic mutation accumulation in fibroblasts of DNA mismatch repair deficient *Msh2^-/-^* (*26*). Thus far, there is very little information about somatic SVs, which cannot be detected accurately, either using single-cell or single-molecule methods(*69*). While their relative frequency is much lower than that of SNVs, SVs can have a major phenotypic impact, as indicated by germline SVs in human diseases(*70*). SVs are much more likely to adversely affect cell function than SNVs, but unless they result in cell death, a mechanism to select against them in non-dividing cells *in vivo* during aging is difficult to imagine.

Future efforts should extend the framework presented in this study to additional cell types, especially quiescent populations, and incorporate methods that capture structural variation. Integrating mutational landscapes with functional readouts *in vivo* will be crucial for determining how the balance between information loss and negative selection shapes organismal aging and disease.

## Methods

### Cell culture and treatment

Human IMR-90 primary fibroblasts were obtained from the American Tissue Culture Collection (ATCC, Lot 70020198). They were cultured in DMEM (Dulbecco’s Modified Eagle Medium, from ATCC) supplemented with 10% fetal bovine serum (FBS, Gibco/Life Technologies) and 1% Penicillin-Streptomycin solution (Gibco/Life Technologies) at 3% oxygen and 37°C. Cells obtained from ATCC were at passage 11 at arrival and were cultured for an additional 5 passages before the start of the experiment. At the beginning of each cycle, 1*10^6^ human primary lung fibroblasts (IMR-90) cells were counted using a Countess 3 (Fisher Scientific), plated, and allowed to expand for 24 hours. Cells were then treated with 50 µg/mL N-ethyl-N-nitrosourea (ENU) and allowed to recover for 3 days. Cells were then collected and assayed for cell number and viability for the 3-day time point. Of the collected cells, 1*10^6^ cells were re-plated and then allowed another 4 days to recover and expand until they were collected to assess cell number and viability for the 7-day time point. This procedure was repeated for 9 consecutive cycles over the course of ∼10 weeks. This experiment was repeated 3 separate times to create 3 independent biological replicates. Cells were isolated at the 7-day time point at cycles 1, 3, 6, and 9 for somatic mutation analysis and at cycles 1 and 9 for transcriptome analysis.

### Cell growth and apoptosis assays

Cells were counted using the ViaCount assay (Cytek) on a Guava EasyCyte flow cytometer to assess cell number and viability at each time point. Apoptosis rates were determined utilizing the Nexin assay (Cytek) on the same machine. Population doublings (PDs) is defined as how many times a population of cells has doubled over a given period of time, was was calculated as:

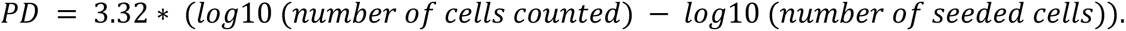

### Bulk mRNA transcriptome analysis

RNA was extracted from frozen cell pellets using the Zymo DNA/RNA extraction kit (Microprep plus D7005) according to the manufacturer’s instructions and quantified using a Qubit kit (Thermo Fisher Scientific). Libraries for mRNA analysis were prepared and sequenced using 150 paired-end mode on an Illumina Novaseq platform by Novogene Corp Inc., CA. Raw sequence reads were processed using the nf-core RNA-Seq pipeline (version 3.1.0) to obtain gene expression levels(*71, 72*). Gene filtering was performed using the zPKM program (Figure S1A)(*73*). Count normalization and differential expression testing was performed using DESeq2 where the design was set to test for the effect of the cycle-condition variable while controlling for the batch variable (Supplementary Note 1)(*74*). Gene set enrichment analysis (GSEA) was performed using the clusterProfiler program (*75*) along with KEGG DNA repair, KEGG apoptosis (*76*) and the SenMayo gene sets(*77*).

### Single-cell isolation, whole-genome amplification, library preparation, and sequencing

Single-cells were isolated using the CellRaft AIR system (Cell Microsystems). The CellRaft array was prepared following the manufacturer’s instruction and washed three times for 3 minutes with warm PBS before addition of cell suspension. 1,000 cells were seeded in 1 mL for one CellRaft array. Cells were allowed to settle for 3-4 hours at 37°C before isolation. Then, the raft was carefully washed with warm medium to remove potential debris and dead cells. Individual rafts containing one cell were deposited in PCR tubes with 2.5 µL of PBS and stored at −80°C until further processing. Single-cell whole-genome amplification and library preparation for WGS was performed as previously described [12]. In brief, single-cell whole genomes were amplified using SCMDA and subject to quality control using a locus dropout test(*78*). Libraries were prepared using a NEBNext Ultra II FS kit (NEB). Library quantity and size were assessed with a Qubit kit and TapeStation, respectively. Libraries were sequenced using the 150 paired-end mode on an Illumina Novoseq platform by Novogene Corp Inc., CA.

### Single-cell whole-genome read alignment and mutation calling

WGS data processing and quality control was performed using as previously described(*24*). Briefly, raw sequencing data underwent quality control using FastQC (*79*) and Trim Galore(*80*), followed by alignment to the human hg19 reference genome using the GATK best practices pipeline(*81*). This included marking adapters, sequence alignment, base quality score recalibration, and variant calling for germline variants using GATK HaplotypeCaller. SCcaller (version 2.0) (*24*)was then employed to call somatic single-nucleotide variations (SNVs) and small insertions and deletions (INDELs), adjusting for allelic amplification bias using known germline variants. Estimated mutation burden was calculated as follows:

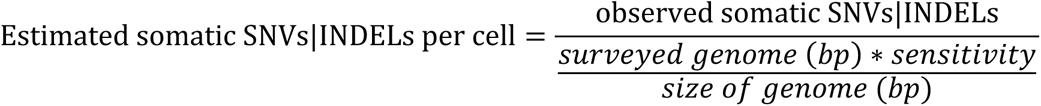

### Mutation filtering and identification of technical artifacts in single-cell whole-genome mutation data

Overlapping SNVs and INDELs were observed amongst the single-cells from batches 2 and 3 (Figures S2D-E; Table S3). Since this was not observed in the first batch, it is unlikely that these mutations are the result of clonal amplification. Mutations that were shared between all cells within a batch were thus considered to be germline and were filtered out prior to downstream analysis (Figures S2F-G).

Technical artifacts inherent to single-cell MDA library preparation steps, such as overheating during cell lysis or whole genome amplification, can be identified by their specific mutational signatures scE and scF(*82*). We checked for the presence of these signatures and found scF to be present in the signature extracted from control cells at cycle 1 (Figure S3A-C). The scF technical artifact is characterized by higher than expected cytosine-to-thymine (C>T) transversions at G<u>C</u>N contexts (Figures SF-G). Thus, control cells from cycle 1 were excluded from the downstream mutational signature analyses.

In an attempt to account for the C>T transition artifacts in control cells at cycle 1, we compared the proportion of the C>T transitions in control cells at cycle 1 to control cells at cycle 9, which do not show any mutational signatures with similarity to the scF sequencing artifact mutational signature (*82*) and have mutation spectra previously observed for IMR90 cells(*83*). We found that this artifact increased the proportion of C>T transitions by 18%±1%. With this, we estimated the amount of artifactual C>T transitions in control cells at cycle 1, subtracted this from the observed mutation burden, and then re-estimated the SNVs per cell (Figure S3D, Table S3).

### Mutation signature analysis

The MutationalPatterns program (*84*) was used to perform mutational spectra and signature analysis. We obtained a diverse set of samples from SomaMutDB (*85*) to use as the background reference mutational signature analysis. These were single-cell whole genome sequencing samples from 10 fibroblast cells(*23*), 199 brain cells(*30*),and 53 lung cells(*29*), where genomic DNA was amplified with either Multiple Displacement Amplification (MDA) or Primary Template-directed Amplification (PTA) (Figure S3E). To determine the number of mutational signatures to be extracted from the data, a non-negative matrix factorization (NMF) rank survey was employed which resulted in 2 factors (Figures S3F). *De novo* signatures were extracted and then compared to those from the COSMIC and SIGNAL databases using cosine similarity(*86, 87*).

### Variant annotation

Observed and expected mutations from each single-cell (or simukation instance of a single-cell in the case of expected mutations) were annotated using the Ensembl Variant Effect Prediction program (version 110.1) and the given ‘Consequences’ annotation was used for downstream analyses(*34*). In the case where mutations were predicted to lead to more than 1 unique consequence (i.e. multiple entries of the same mutation), the highest ranking annotation that occurred in the resulting output was kept which resulted in a 1:1 mapping of mutations and consequences. In the case where mutations occurred in a region that overlapped multiple possible consequences (e.g. splice region variant and intron variant), we classified this mutation to be consistent with the consequence that occurred first in this annotation (e.g. splice region, if multiple annotation was splice region variant and intron variant). For mutations annotated as splice variants (splice_acceptor_variant, splice_donor_variant, splice_donor_5th_base_variant, splice_region_variant,splice_donor_region_variant, splice_polypyrimidine_tract_variant), we classified these mutations as ‘splice.’ Synonymous variants were classified as mutations annotated as synonymous_variant or stop_retained_variant. Non-synonymous variants were classified as mutations annotated as missense_variant, start_lost, stop_gained, or stop_lost.

### Selection analysis of annotated variants

Selection analysis was performed by first simulating 10,000 instances of each single-cell in our dataset to be used as the null expectation (i.e. neutral selection) where mutations are randomly re-distributed throughout the genome. This was done using the SigProfilerSimulator program(*33*), where for each single-cell, observed mutations were randomly re-distributed across the genome while accounting for cell-specific sequencing coverage, mutational signatures, and nucleotide contexts.

Log_2_(observed/expected) ratios (written hereafter as Log_2_(O/E)) were calculated for each single-cell or by group (mutations from all cells within a group are pooled), by first summing the frequency of annotated consequences for observed and expected mutations. The Log_2_(O/E) ratio was then calculated with respect to each simulation instance (resulting in 10,000 Log_2_(O/E) ratios per cell or group) using the following equation:

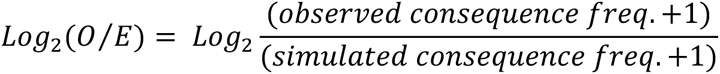

A permutation test was then used to determine the p-value of the observed consequence frequency where the simulated consequence frequency served as the null distribution and is as follows:

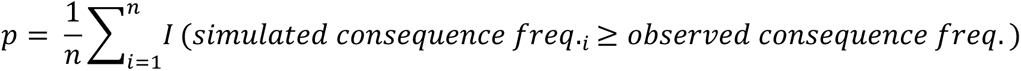

Where *p* is the permutation test p-value, *n* is the number of permutations (i.e. 10,000), *i* is the *i*-th permutation, and *I* is the indicator function, which equals 1 if the condition inside is true, and 0 otherwise.

We performed a power analysis to determine how many observed variants are required to obtain reliable statistical results at varying Log_2_(O/E) ratios. To do this, we used the mutation data from ENU-treated cells to obtain the relationship between the standard deviation in the number of expected variants given the number of observed variants for common types of variants that were identified. We fit a gamma GLM with a log link to model standard deviation as a function of log-transformed observed variant count. The model showed excellent fit (residual deviance = 3.19 on 144 df vs. null deviance = 147.37 on 145 df; dispersion = 0.020), with log-transformed observed variant count as a highly significant predictor irrespective of the variant type (p < 2 × 10⁻¹⁶). With this, we then calculated Log_2_(O/E) ratios by simulating 10,000 instances of expected variant counts while varying the observed variant counts from 1-1,000. We then varied the Log_2_(O/E) ratios from 0-1 by multiplying the observed variant count by a scaling factor before calculating the Log_2_(O/E) ratio. Finally, we calculated p-values for the Log_2_(O/E) ratios using a permutation test as described previously.

The power analysis demonstrated how varying the observed variant count affects the p-value of the Log_2_(O/E) ratio at different levels. Importantly, this analysis provided us with estimates of how many observed variants were required to achieve sufficient power (p-value<0.05) (i.e. obtain reliable statistical results) given a specific Log_2_(O/E) (Figure S4B, solid line). This threshold was then applied to the Log_2_(O/E) results, at both the levels of variant types and pathways, to determine which of these were amenable to further statistical testing where the results of this can be seen in the figure legends (e.g. Figure 5B; Table S5). For the Log_2_(O/E) non-synonymous/synonymous calculation, we determined if these were amenable to further statistical testing if either the non-synonymous or synonymous Log_2_(O/E) results in each group reached the established observed variant threshold (Table S5).

When assessing for mutation selection, it is also necessary to account for the cell-intrinsic bias in mutation frequency due to DNA repair mechanisms (i.e. transcription coupled repair), which is known to reduce mutation frequency in transcribed regions and is agnostic to damage that would result in synonymous or non-synonymous variants(*88*). This bias in mutation frequency was indeed observed in our data and consistent with a depletion of mutations due to DNA repair (Figure 5H). Thus, we normalized the Log_2_(O/E) ratio of coding variants (e.g. synonymous, missense, etc.) and non-coding variants (5’/3’ UTR and splice region) by the Log_2_(O/E) ratio of mutations which occurred within intronic regions and not in splice regions. We chose to use this normalization because the occurrence of both exonic and intronic mutations is dependent on transcription coupled repair (*37*) however, since intronic mutations do not alter the translated protein (with the exception of splice site variants), they are not subject to selective mechanisms. In addition, the majority of variants in our data occur within introns (∼46.7%, Figure 5G), allowing us to make more confident Log_2_(O/E) ratio for these mutations as compared to using the Log_2_(O/E) ratio of synonymous mutations which only made up ∼0.23% of all variants. A comparison of unnormalized and normalized Log_2_(O/E) ratio for non-synonymous and synonymous variants is shown for example by comparing Figures 5B to S4C, where the effect of normalization can be seen to slightly increase the Log_2_(O/E) ratio.

To estimate the number of variants that are protected against resulting from negative selection, we subtracted the average number of variants from simulated samples from the number of variants in the respective observed sample.

### dN/dS analysis

For an alternative measure of selection, we employed the dndscv R package(*39*), which utilizes a trinucleotide context-dependent substitution model to estimate dN/dS ratios. The analysis was performed using the hg19 reference genome. We used default settings with the exception of disabling the filtering of hypermutated samples and genes (max_coding_muts_per_sample = Inf and max_muts_per_gene_per_sample = Inf) to retain all mutation data in the cohort.

### Selection analysis of gene sets and pathways

Non-expressed and expressed genes were determined using bulk mRNA transcriptome data from cycle 9 control and ENU-treated groups using the zPKM program (Figure S2A; Table S7)(*73*). Nonessential and essential control gene sets were obtained from the DepMap database (depmap.org)(*89*), where the nonessential gene set was the negative controls from Hart *et al* (2015) and the essential gene set was the intersection of the the essential gene sets resulting from the Hart *et al* (2015) and Blomen *et al* (2014) studies(*48, 90*). These gene sets were additionally filtered for expressed genes by overlapping them with the list of expressed genes we determined previously (Table S7). Haplosufficient and haploinsufficient gene sets were obtained from the ExAC database where haplosufficient genes were defined as pLI < 0.1 and haploinsufficient genes were defined as pLI > 0.9(*35*). These gene sets were also filtered for expressed genes in the same manner as essential genes (Table S7). Pathway level gene sets were obtained by performing over representation analysis for gene ontology biological processes using the clusterProfiler program (*75*) on the list of expressed genes that determined previously (Table S8). For all gene sets, genomic coordinates of exons, introns, promoters (5KB upstream of transcription start site), and annotated enhancers (*44*) were retrieved using the GenomicRanges (version 1.60.0) software(*91*).

For each gene set, two Log_2_(O/E) ratios were calculated. One for the version of the gene set where the intronic regions were included, and the other for the version of the gene set intronic regions were excluded. To account for DNA repair (as previously described for Log_2_(O/E) calculation of coding and non-coding variants), the Log_2_(O/E) ratios of each gene set was normalized by subtracting Log_2_(O/E) ratio of the version that included introns from the version that excluded introns. A permutation test was then used to estimate p-values, as described above, and were adjusted for multiple hypothesis testing using the Benjamini-Hochberg method.

To serve as a negative control for the pathway analysis, for each enriched biological process gene set, we randomly shuffled the genes between all other identified biological processes gene sets and created 10,000 randomly permuted gene sets. Average Log_2_(O/E) ratios were then calculated for the 10,000 permutations of each gene set followed by p-value estimation, as described above.

## Data and code availability

Raw sequencing data has been deposited in the NCBI Sequence Read Archive under BioProject accession PRJNA1129131. Processed gene expression data has been deposited in the NCBI Gene Expression Omnibus database under accession GSE271867. Processed variant call files have been deposited in the NCBI dbSNP database and will be made available upon acceptance. Scripts used for processing and analyzing data have been deposited at:

figshare.com/articles/software/Negative_selection_allows_human_primary_fibroblasts_to_tolera te_high_somatic_mutation_loads_induced_by_N-ethyl-N-nitrosourea/26110453

## Competing interests

M.L., A.Y.M., X.D., and J.V. are co-founders and shareholders of SingulOmics Corp. A.Y.M., and J.V. are co-founders of MutagenTech Inc. Other authors declare no conflict of interest.

## Acknowledgements

We would like to acknowledge the members of the Vijg and Sidoli labs for helpful discussions during the preparation of the manuscript.

## Author Contributions

A.Y.M., and J.V. conceived this study and designed the experiments. J.H., M.S., and A.Y.M performed the experiments. R.C., J.H., and S.S. analyzed the data. X.D. provided guidance on the data analysis. R.C., J.H., and J.V. wrote the manuscript.

## Funding

This study was supported by National Institute of Health grants: T32GM007491 (R.C.) T32AG023475 (R.C.), U19AG056278 (J.V.), U01HL145560 (J.V.), U01ES029519 (J.V.), P01AG017242 (J.V.), P01AG047200 (J.V.), RF1AG068908 (J.V.), and P30AG038072 (J.V.). Other funding sources: US Department of Defense grant BC180689P1 (J.V.), Glenn Foundation for Medical Research Postdoctoral Fellowships in Aging Research (J.H.), The Michael J. Fox Foundation (J.V.), and The Paul F. Glenn Center for the Biology of Human Aging at the Albert Einstein College of Medicine (J.V.).

## Notes

### Summary of Updates

Clarified conceptual and methodological distinctions between observed, estimated, and expected mutations; expanded and refined our analyses of negative selection in both coding and non-coding regions; and added new explanations, controls, and supplementary materials.

