## Supplementary Materials for "Evidence for negative selection against somatic mutations in fibroblasts with high N-ethyl-N-nitrosourea-induced mutation load"

### Contents:


Supplementary Note 1
Supplementary Note 2
Supplementary Note 3
Supplementary Note 4
Supplementary Note 5

| | |
|---|---|
| Supplementary Figure 1: | Transcriptome Analysis Quality Control and Extended Analysis. Related to Figure 2. |
| Supplementary Figure 2: | Single-Cell Whole Genome Analysis Quality Control and Observed Mutational Burden. Related to Figure 3. |
| Supplementary Figure 3: | Extended analysis of Mutational Spectra and Signatures. Related to Figure 4. |
| Supplementary Figure 4: | Extended Analysis of Negative Selection Against Damaging Coding and Non-Coding Variants in ENU-treated Cells. Related to Figure 5. |
| Supplementary Figure 5: | dN/dS analysis of selection pressure for damaging coding variants. Related to Figure 5. |
| Supplementary Figure 6: | Extended Selection Pressure Analysis of Mutation Accumulation at the Pathway Level. Related to Figure 6. |
| Supplementary Figure 7: | Extended Analysis of Negative Control to Validate Negative Selection of Functional Pathways. Related to Figure 6. |

| | |
|---|---|
| Table S1: | Sample table for bulk mRNA-sequencing experiment. Related to Figures 2 and S1 |
| Table S2: | Differential gene expression results from bulk mRNA-sequencing experiment. Related to Figures 2 and S1. |
| Table S3: | Sample table for single-cell whole genome sequencing experiment with metrics related to mutation calling and mutation burden. Related to Figures 3 and S2. |
| Table S4: | Results of selection analysis ($Log_2$(observed/expected)) for coding and non-coding variants. Related to Figures 5 and S4. |
| Table S5: | Variant effect prediction annotation of all mutations. |
| Table S6. | Results of dndscv for cancer driver gene mutations in ENU-treated cells. |
| Table S7: | List of Entrez IDs for Non-expressed/Expressed, Nonessential/Essential, and Haplosufficient/Haploinsufficient gene sets. Related to Figures 6 and S5. |

| | |
|---|---|
| Table S8: | Gene set enrichment analysis results for gene ontology biological process of expressed genes. Related to Figures 6 and S5. |
| Table S9: | Results of selection analysis ($Log_2$(observed/expected)) for GO biological processes. Related to Figures 6 and S5. |
| Table S10. | Catalogue of all mutations identified in control and ENU-treated cells. Related to Figures 3-6. |

### Supplementary Notes

Supplementary Note 1

Strong batch effects were found to be present during the analysis of the bulk mRNA-Seq data (Figures S1G-I), which could mask the effects of interest when comparing the different groups. While the batch effect was corrected for during the differential expression (DE) analysis by accounting for the batch variable in the generalized linear model utilized by DESeq2 (Figures S1J-K), we also performed the DE analysis of ENU Cycle 9 vs Control Cycle 9 after removing samples from the Control Cycle 1 group. After batch correction, separation between the groups could be seen from principal component analysis (Figure S1L). However, a very small number of genes were differentially expressed, validating our findings then including the Control Cycle 3 group (Figure S1M).

Supplementary Note 2

Though relatively few INDELs were observed (369 total), the mutational spectra in ENU-treated cells were characterized by an increase in thymine INDELs as well as INDELs with a 5+bp deletion and microhomology relative to the controls (Figure S3H). Moreover, a single *de novo* INDEL mutational signature could be extracted, which weakly resembled COSMIC ID1 and ID2 (cosine similarity 0.59 & 0.54, respectively). Both have been suggested to be due to slippage of the template strand during DNA replication in addition to having “clock-like” behavior.

Supplementary Note 3

To test if the negative selection finding could be recapitulated with other software that calculates selection pressure, we used the dNdScv software with the default parameters and as input we used the mutations from the control and ENU-treated groups (Methods). This resulted in a global dN/dS ratio of ~1 for all mutation types in all the ENU-treated groups, indicative of neutral selection (Figure S5A). As these results conflicted with our own results using the O/E method, we sought to determine what the differences were between the two methods and if either of these was better suited to analyzing our single-cell whole genome sequencing (SCWGS ) data. We compared the counts of observed and expected for synonymous (Figures S5B-F) and missense variants (Figures S5G-K) between the methods, which led us to identify the main discrepancy: the approaches used to estimate the expected number of variants. The dndscv approach calculates expected variants per gene based on probabilistic models of mutation rates and sequence contexts (*35*). In contrast, the O/E method uses SigProfilerSimulator to empirically estimate expected variants by simulating mutations randomly across the genome while accounting for coverage, sequence context, and mutational signatures (*32*). Crucially, the dndscv model makes assumptions about uniform genomic coverage and high mutation counts, conditions violated in our SCWGS data, as it is characterized by uneven coverage and relatively few coding mutations per cell (Figures S2A and 5A). Finally, an argument against the null model generated by SigProfilerSimulator in the O/E method (i.e. the expected mutations) is that due to its simplicity, it might not accurately reflect neutral selection as one would expect. However, when simulating 1,000 instances of mutations using SigProfilerSimulator (given observed mutations from the control and ENU groups) and then using the results of this as input to the dndscv program, this produced a global dN/dS ratio of ~1 for all variants, as would be expected (Figure S5L). This means that the null model generated by SigProfilerSimulator accurately models neutral selection pressure, according to the dndscv program. In summary, while dN/dS is a powerful metric for highly mutated cancer genomes, it lacks sensitivity to detect negative selection in scenarios like ours with limited coding mutations and variable coverage across the genome, which is inherent to SCWGS. We conclude that empirically estimating expected variants via simulations, as done in our O/E analysis, is better suited for such situations.

<u>Supplementary Note 4</u>

Out of the 37 detected stop gain variants, 4 of these were in 4 different cancer driver genes and had dN/dS ratios from ~70-1,000 when analyzing selection using dndscv, though they were not significant after multiple hypothesis correction. However, these particular variants could possibly explain the positively trending $Log_2$(O/E) ratio seen for stop gain variants in the ENU-treated cells (Figure 5E). These variants were found in *ABL1* and *CDC73*, which are tumor suppressors while *CREB3L2* and *COL1A1* are oncogenes (Table S6). Thus, it may be possible that stop gain variants in ABL1 and CDC73 may have provided a slight growth advantage that led it to being picked up in our analysis. However, repeated passaging during ENU treatment creates a strong bottleneck that limits clonal expansion. While 8 shared SNVs were detected between 2 cells, the presence of ~9,000 unique SNVs per cell indicates these arose independently, arguing against large clonal expansion (Figure S2F). It will be interesting for further studies which analyze larger numbers of somatic mutations in normal cells to see if this may be true.

In an attempt to gain insight in selection pressure at the gene level, we mapped the non-synonymous variants to the COSMIC Cancer Gene Census gene set and identified a total of 75 cancer driver genes containing nonsynonymous variants (Methods). However, low variant counts of ~1 per gene on average essentially constrained meaningful conclusions when assessing selection pressure with either the O/E or dNdScv methods (Table S6).

<u>Supplementary Note 5</u>

Given that there are enough observed variants for sufficient statistical power and after accounting for transcription coupled repair (Methods), negative $Log_2$O/E ratios suggest negative selection because the frequency of observed variants is less than what would be expected according to the null model. In contrast, positive $Log_2$O/E ratios do not necessarily positive selection. This is because in the calculation of the $Log_2$O/E ratio, while mutations are randomly re-distributed in the simulated samples according to the null model, the total number of variants remains the same between observed and simulated samples. This creates the potential for a see-saw effect as a depletion of mutations in a large regions (i.e., introns, which make up ~25% of the genome) must be balanced out. The consequence of this may be an inflated estimate of the $Log_2$O/E ratio for other large regions (i.e., intergenic regions, which make up ~50% of the genome) (Figure 5I), which warrants caution for the interpretation of these results. Despite this potential limitation of the O/E method, we do believe that the resulting positive $Log_2O/E$ ratios for intergenic regions is not the result of such technical artifact, as this is supported by previous evidence that intergenic regions lack DNA repair activity and are under neutral selection (*36, 37, 88*).

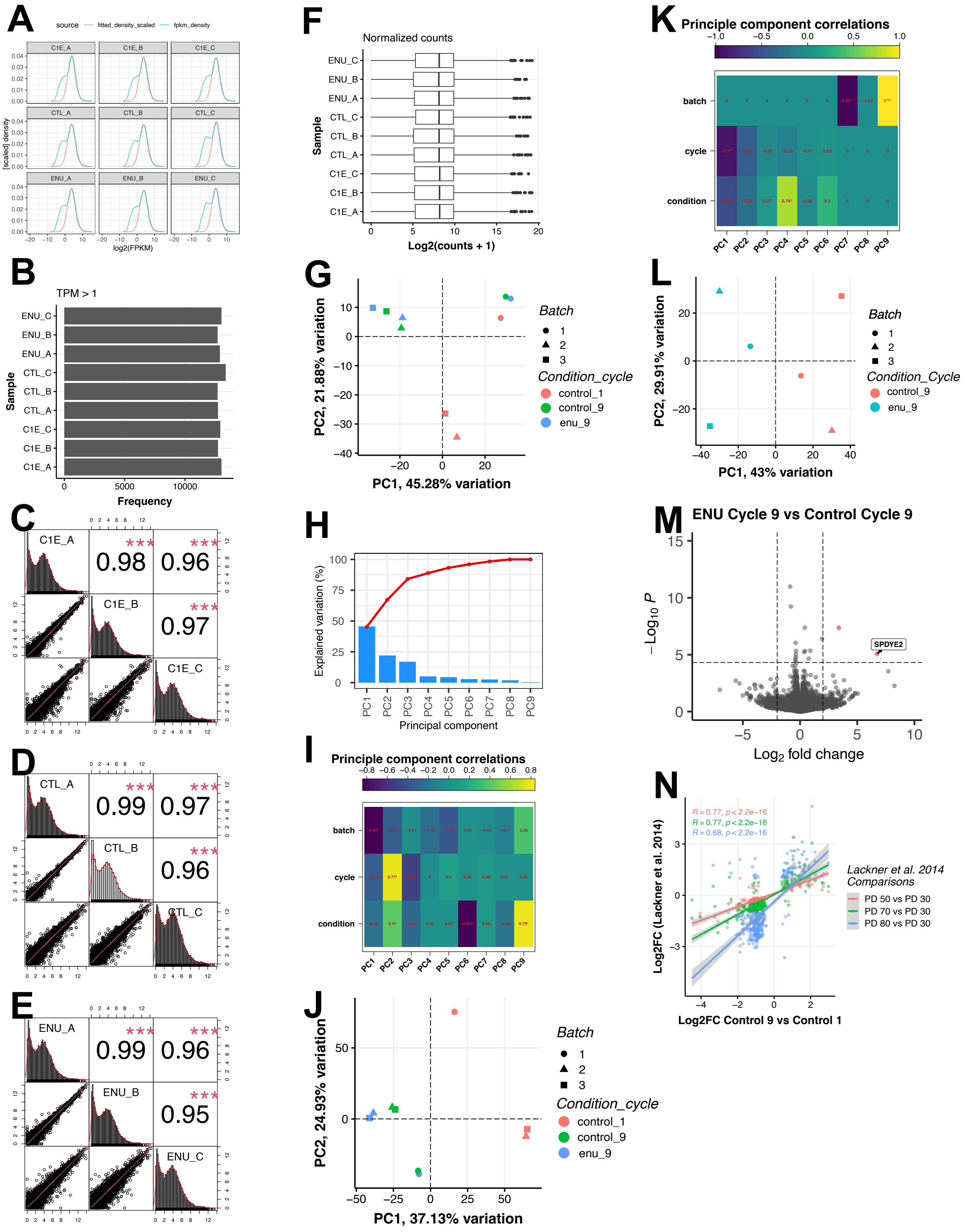

A
source
fitted_density_scaled
fpkm_density
[scaled] density
log2(FPKM)
C1E_A
C1E_B
C1E_C
CTL_A
CTL_B
CTL_C
ENU_A
ENU_B
ENU_C
B
TPM > 1
Sample
Frequency
C
0.98
0.96
0.97
D
0.99
0.97
0.96
E
0.99
0.96
0.95
F
Normalized counts
Sample
Log2(counts + 1)
G
PC1, 45.28% variation
PC2, 21.88% variation
Batch
Condition_cycle
control_1
control_9
enu_9
H
Explained variation (%)
Principal component
I
Principle component correlations
batch
cycle
condition
J
PC1, 37.13% variation
PC2, 24.93% variation
Batch
Condition_cycle
control_1
control_9
enu_9
K
Principle component correlations
batch
cycle
condition
L
PC1, 43% variation
PC2, 29.91% variation
Batch
Condition_Cycle
control_9
enu_9
M
ENU Cycle 9 vs Control Cycle 9
SPDYE2
$-Log_{10}$ P
$Log_2$ fold change
N
R = 0.77, p < 2.2e−16
R = 0.77, p < 2.2e−16
R = 0.68, p < 2.2e−16
Log2FC (Lackner et al. 2014)
Log2FC Control 9 vs Control 1
Lackner et al. 2014 Comparisons
PD 50 vs PD 30
PD 70 vs PD 30
PD 80 vs PD 30

**Supplementary Figure 1. Transcriptome Analysis Quality Control and Extended Analysis. Related to Figure 2.**

**(A)** Gene abundance distribution of each sample showing unfiltered distribution (blue) and the Gaussian fit to the $Log_2$(FPKM) transcript abundances (red), which represents the filtering threshold used to identify expressed genes used in downstream analyses. **(B)** Number of genes detected at TPM>1. **(C-E)** Correlation of $Log_2$(TPM) data between biological replicates in control cycle 1 (C), control cycle 9 (D), and ENU-treated cycle 9 (E) groups. Statistics: n=16,133; Pearson R correlation coefficient; p-value were estimated using a t-test; p-value legend: ***: p <= 0.001. **(F)** Distribution of DESeq2 normalized gene abundances. **(G)** Principal component analysis (PCA) of all samples. Note the apparent batch effect. **(H)** Percent of variance explained by each PC in (G). **(I)** Correlation of each PC to experimental variables, showing the batch effect that is captured by the first PC. Statistics: n=3; Pearson R correlation coefficient; p-value were estimated using a t-test. **: p <= 0.01. **(J)** PCA of all samples after adjusting for the batch variable (Methods). **(K)** Correlation of each PC to experimental variables after batch correction, showing a lack of correlation of the batch variable with the first 6 PCs. Statistics: Same as in (I). **(L)** PCA of samples only from cycle 9 after adjusting for the batch variable (Methods). See Supplemental Note 1 for details on the purpose of this analysis. **(M)** Differential gene expression results of ENU-treated cycle 9 vs. control cycle 9 when only analyzing cycle 9 samples after adjusting for the batch variable (as shown in L). Red points are $Log_2$ fold change>2 and adjusted p-value<0.05. Statistics: n=3; DGE results obtained using Wald test followed by Benjamini-Hochberg correction for multiple hypothesis testing.

193

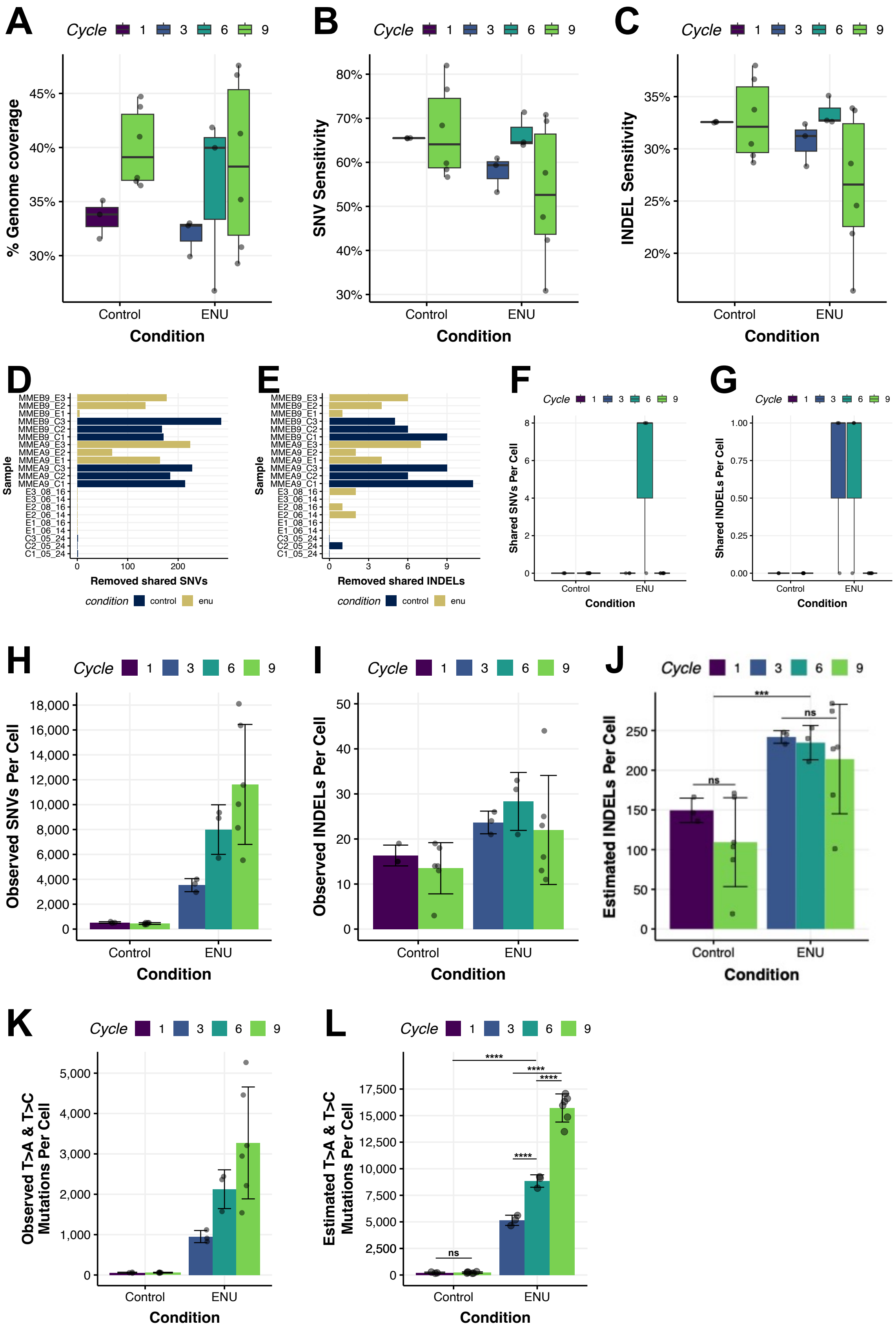

A
Cycle 1 3 6 9
% Genome coverage
30%
35%
40%
45%
Control
ENU
Condition
B
SNV Sensitivity
30%
40%
50%
60%
70%
80%
C
INDEL Sensitivity
20%
25%
30%
35%
D
Sample
MMEB9_E3
MMEB9_E2
MMEB9_E1
MMEB9_C3
MMEB9_C2
MMEB9_C1
MMEA9_E3
MMEA9_E2
MMEA9_E1
MMEA9_C3
MMEA9_C2
MMEA9_C1
E3_08_16
E3_06_14
E2_08_16
E2_06_14
E1_08_16
E1_06_14
C3_05_24
C2_05_24
C1_05_24
Removed shared SNVs
condition control enu
E
Removed shared INDELs
F
Shared SNVs Per Cell
G
Shared INDELs Per Cell
H
Observed SNVs Per Cell
I
Observed INDELs Per Cell
J
Estimated INDELs Per Cell
ns
***
K
Observed T>A & T>C Mutations Per Cell
L
Estimated T>A & T>C Mutations Per Cell
****

**Supplementary Figure 2. Single-Cell Whole Genome Analysis Quality Control and Observed Mutational Burden. Related to Figure 3.**

**(A)** Percentage of genome covered at 20X read depth per cell. **(B-C)** Sensitivity of SNV (B) and INDEL (C) mutation calling (Methods). **(D-E)** Number of observed SNVs (D) and INDELs (E) removed from cells collected in the same cycle and batch that shared SNVs and INDELs (Methods). **(F-G)** Number of observed shared SNVs and INDELs amongst all cells after filtering for mutations shared within each cycle and batch (Methods). **(H-I)** Observed SNVs (H), INDELs (I) per cell. **(J)** Estimated burden of small insertions/deletions and (INDELs) per cell corrected for genome coverage and sensitivity. Statistics: Data represent mean ± S.D; p-values of comparisons between cycles within the same condition were estimated with a two-sided linear model with estimated marginal means, Tukey's HSD. p-values of comparisons of control and ENU-treated groups were estimated using a two-way ANOVA and are shown at top of plots; p-value legend ***: p <= 0.001. **(K)** Observed T>A & T>C SNVs, attributed to ENU, per cell. Statistics: Same as in (H) **(L)** Estimated T>A & T>C SNVs, attributed to ENU, per cell. Statistics: Same as in (J). Statistics for all panels: n=3 cells for cycles 1, 3, and 6; n=6 cells for cycle 9. Data represent mean ± S.D.

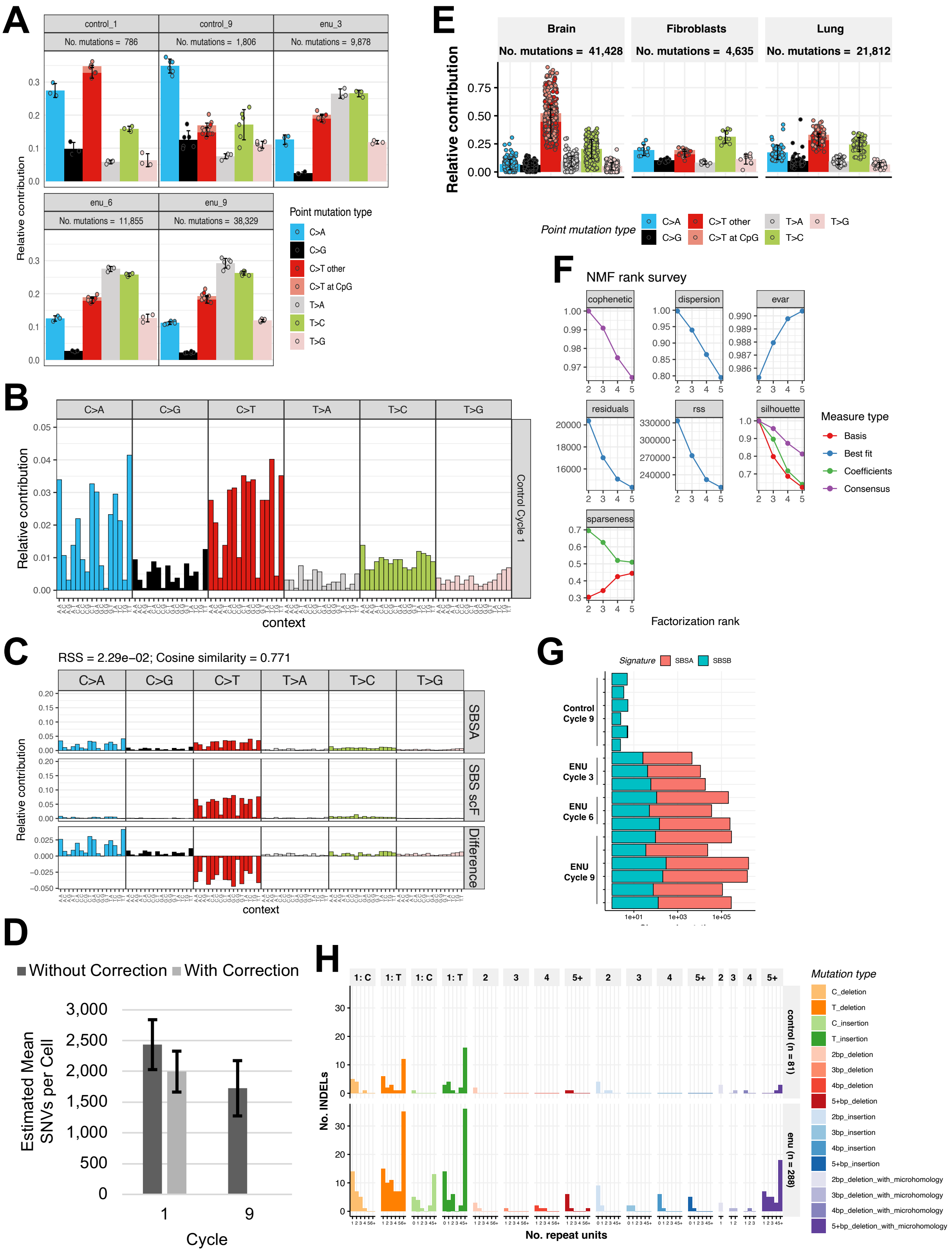
A
control_1
No. mutations = 786
control_9
No. mutations = 1,806
enu_3
No. mutations = 9,878
enu_6
No. mutations = 11,855
enu_9
No. mutations = 38,329
Relative contribution
Point mutation type
C>A
C>G
C>T other
C>T at CpG
T>A
T>C
T>G
B
C>A
C>G
C>T
T>A
T>C
T>G
Relative contribution
Control Cycle 1
context
C
RSS = 2.29e−02; Cosine similarity = 0.771
SBSA
SBS scF
Difference
Relative contribution
context
D
Without Correction
With Correction
Estimated Mean SNVs per Cell
Cycle
E
Brain
No. mutations = 41,428
Fibroblasts
No. mutations = 4,635
Lung
No. mutations = 21,812
Relative contribution
Point mutation type
F
NMF rank survey
cophenetic
dispersion
evar
residuals
rss
silhouette
sparseness
Measure type
Basis
Best fit
Coefficients
Consensus
Factorization rank
G
Signature
SBSA
SBSB
Control Cycle 9
ENU Cycle 3
ENU Cycle 6
ENU Cycle 9
H
No. INDELs
No. repeat units
control (n = 81)
enu (n = 288)
Mutation type
C_deletion
T_deletion
C_insertion
T_insertion
2bp_deletion
3bp_deletion
4bp_deletion
5+bp_deletion
2bp_insertion
3bp_insertion
4bp_insertion
5+bp_insertion
2bp_deletion_with_microhomology
3bp_deletion_with_microhomology
4bp_deletion_with_microhomology
5+bp_deletion_with_microhomology

**Supplementary Figure 3. Extended analysis of Mutational Spectra and Signatures. Related to Figure 4.**

**(A)** Mutational spectra of the relative contribution of different types of observed SNVs per cell separated by cycle and condition. Note the abnormal increase of C>T transition indicative of a technical artifact in control cycle 1 group, which was removed from downstream signature analysis (Methods). Related to Figure 4A. Statistics: Data represent mean±S.D.; n=3 cells for cycles 1, 3, and 6; n=6 cells for cycle 9. **(B)** Mutational profile of control cycle 1 group, showing higher than expected cytosine-to-thymine (C>T) transversions at G<u>C</u>N contexts. Statistics: n=3. See Methods for details on the exclusion of this group from this analysis. **(C)** Comparison of the only mutational signature extracted from the control cycle 1 group, SBSA, to a known single cell library preparation artifact SBS scF. See Methods for details on the exclusion of this group from this analysis. Statistics: n=3; cosine similarity=0.771, residual sum of squares=$2.29*10^{-2}$ . **(D)** Estimated burden of somatic single nucleotide variants (SNVs) per cell in the control groups corrected for genome coverage, sensitivity, and the scF sequencing artifact mutational signature (B-C). This artifact increased the proportion of C>T transitions by 18%±1%. With this, we estimated the amount of artifactual C>T transitions in control cells at cycle 1, subtracted this from the observed mutation burden, and then re-estimated the SNVs per cell. Statistics: n=3 cells for cycle 1; n=6 cells for cycle 9; data represent mean ± S.D. **(E)** Mutational spectra of the relative contribution of different types of observed SNVs per cell separated by study representing brain (*27*), fibroblast (*21*), and lung (*26*) tissues. These additional samples were included to be used as background spectra for signature extraction. Statistics: n=199 brain cells; n=10 fibroblasts cells; n=53 lung cells; data represent mean ± S.D. **(F)** Results of determining optimal factorization rank for SNV signature extraction using non-negative matrix factorization, where a rank of 2 was chosen to extract 2 mutational signatures from the mutation data. Related to Figure 4B. **(G)** Absolute contribution of *de novo* mutational signatures SBSA and SBSB to individual cells (shown as rows) grouped by condition and cycle. Related to Figure 4B. **(H)** Small insertion and deletion (INDEL) mutation spectra grouped by condition. See Supplemental Note 2 for detailed description of each spectrum. Statistics: n=6 cells for control group; n=12 cells for ENU-treated group.

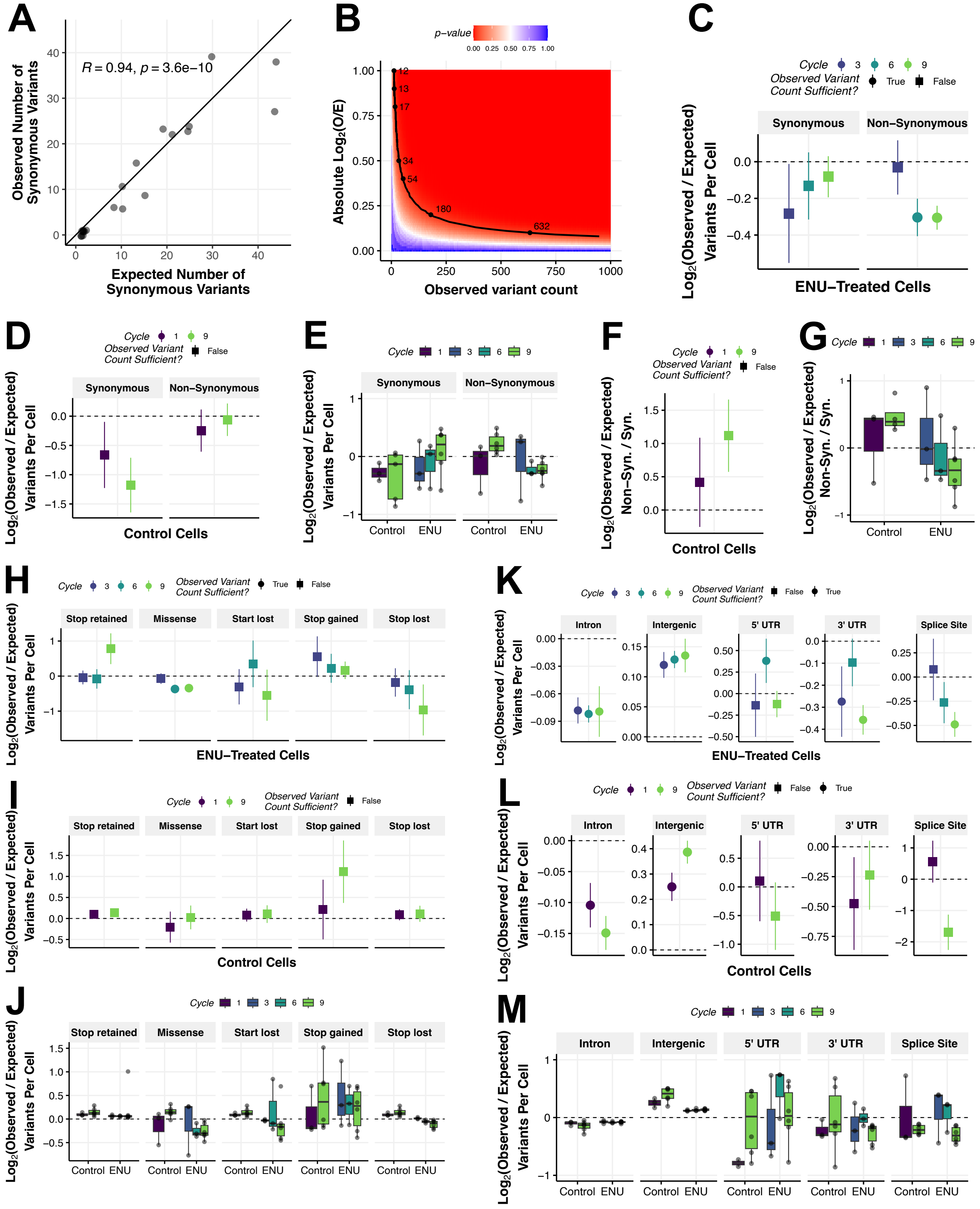
A
R = 0.94, p = 3.6e−10
Observed Number of Synonymous Variants
Expected Number of Synonymous Variants
B
p−value
Absolute Log2(O/E)
Observed variant count
C
Cycle
Observed Variant Count Sufficient?
True
False
Synonymous
Non−Synonymous
Log2(Observed / Expected) Variants Per Cell
ENU−Treated Cells
D
Control Cells
E
Control
ENU
F
Log2(Observed / Expected) Non−Syn. / Syn.
G
H
Stop retained
Missense
Start lost
Stop gained
Stop lost
I
J
K
Intron
Intergenic
5' UTR
3' UTR
Splice Site
L
M

**Supplementary Figure 4. Extended Analysis of Negative Selection Against Damaging Coding and Non-Coding Variants in ENU-treated Cells. Related to Figure 5.**

**(A)** Correlation of observed and expected (simulated) synonymous variants for each cell. The black line indicates a perfect correlation (slope=1). Shown at top is the fit of a linear equation. Statistics: n=21; fitted line represents mean ± S.D.; Pearson R correlation coefficient and p-value of the correlation was estimated using a T-test. **(B)** P-value as a function of the observed variant count and the $Log_2(O/E)$ to determine the number of observed variants given the $Log_2(O/E)$ that is needed to reliably interpret the resulting statistical test (Methods). Related to Figures 5 and 6. **(C)** Unnormalized $Log_2(O/E)$ ratio of synonymous and non-synonymous variants for ENU-treated cells, related to Figure 5B. **(D)** Normalized $Log_2(O/E)$ ratio of synonymous and non-synonymous variants for control cells, related to Figure 5B. **(E)** Normalized $Log_2(O/E)$ ratio of synonymous and non-synonymous variants per cell, related to Figure 5B. **(F)** Normalized $Log_2(O/E)$ ratio of the non-synonymous / synonymous (N/S) ratio for control cells, related to Figure 5D. **(G)** Normalized $Log_2(O/E)$ ratio of the non-synonymous / synonymous (N/S) ratio per cell for control and ENU-treated cells, related to Figure 5D. **(H)** Unnormalized $Log_2(O/E)$ ratio of coding variants for ENU-treated cells, related to Figure 5F. **(I)** Normalized $Log_2(O/E)$ ratio of coding variants for control cells, related to Figure 5F. **(J)** Normalized $Log_2(O/E)$ ratio of coding variants per cell, related to Figure 5F. **(K)** Unnormalized $Log_2(O/E)$ ratio of non-coding variants for ENU-treated cells, related to Figure 5H. **(L)** Normalized $Log_2(O/E)$ ratio of non-coding variants for control cells, related to Figure 5H. **(M)** Normalized $Log_2(O/E)$ ratio of non-coding variants per cell, related to Figure 5H. Statistics for all panels: n=3 cells for cycles 1, 3, and 6; n=6 cells for cycle 9; Data represent mean ± S.D.

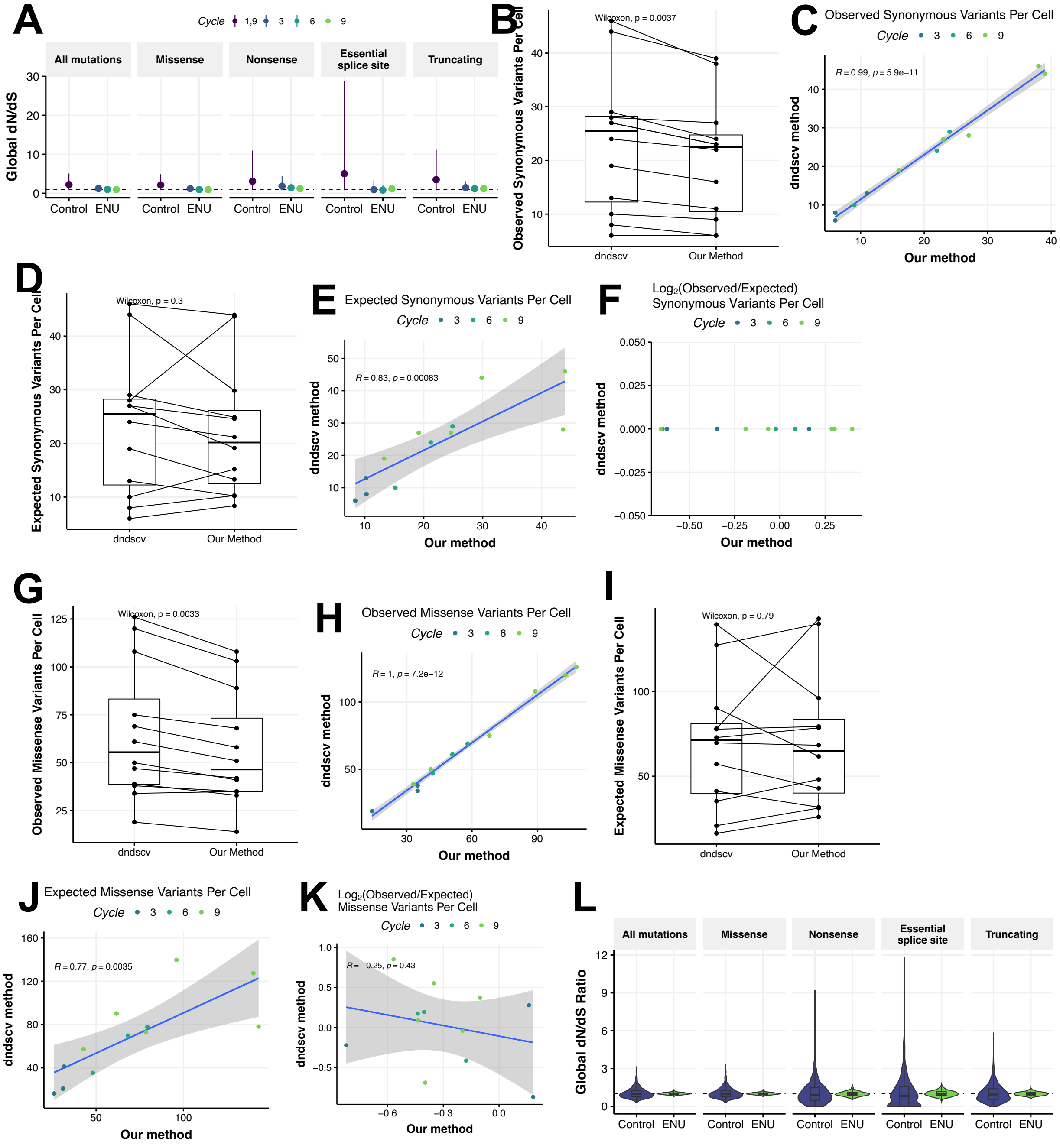
A
Cycle 1,9 3 6 9
All mutations
Missense
Nonsense
Essential splice site
Truncating
Global dN/dS
Control ENU
B
Wilcoxon, p = 0.0037
Observed Synonymous Variants Per Cell
dndscv Our Method
C
Observed Synonymous Variants Per Cell
Cycle 3 6 9
R = 0.99, p = 5.9e−11
dndscv method
Our method
D
Wilcoxon, p = 0.3
Expected Synonymous Variants Per Cell
dndscv Our Method
E
Expected Synonymous Variants Per Cell
Cycle 3 6 9
R = 0.83, p = 0.00083
dndscv method
Our method
F
Log2(Observed/Expected)
Synonymous Variants Per Cell
Cycle 3 6 9
dndscv method
Our method
G
Wilcoxon, p = 0.0033
Observed Missense Variants Per Cell
dndscv Our Method
H
Observed Missense Variants Per Cell
Cycle 3 6 9
R = 1, p = 7.2e−12
dndscv method
Our method
I
Wilcoxon, p = 0.79
Expected Missense Variants Per Cell
dndscv Our Method
J
Expected Missense Variants Per Cell
Cycle 3 6 9
R = 0.77, p = 0.0035
dndscv method
Our method
K
Log2(Observed/Expected)
Missense Variants Per Cell
Cycle 3 6 9
R = −0.25, p = 0.43
dndscv method
Our method
L
All mutations
Missense
Nonsense
Essential splice site
Truncating
Global dN/dS Ratio
Control ENU

**Supplementary Figure 5. dN/dS analysis of selection pressure for damaging coding variants. Related to Figure 5.**
**(A)** Global dN/dS ratios across various SNV types using observed SNVs as input. Control cells from cycle 1 and 9 are combined due to low total number of mutations. Statistics: n=3 cells for cycles 1, 3, and 6; n=6 cells for cycle 9; data represent mean ± S.E. **(B)** Difference in the number of observed synonymous variants either determined by dNdScv software or our annotation approach using VEP (Methods). Statistics: n=18 cells; p-value estimated using Wilcoxon rank-sum test. **(C)** Same as in (B), but plotted as a scatter. Statistics: n=18 cells; fitted line represents mean ± S.D.; Pearson R correlation coefficient and p-value of the correlation was estimated using a T-test. **(D)** Difference in the estimated number of expected synonymous variants either determined by dNdScv software or our simulation approach using SigProfilerSimulator (Methods). Statistics: Same as in (B). **(E)** Same as in (D), but plotted as a scatter. Statistics: Same as in (C). **(F)** Correlation of synonymous variant $Log_2$(O/E) ratio either determined by dNdScv software (using the observed and expected frequency outputs) or our approach (Methods). Statistics: Same as in (C). **(G-K)** Same as in (B-F), but for missense variants. **(L)** Global dN/dS ratios across various SNV types using expected SNVs as input. Statistics: Same as in (A).

283

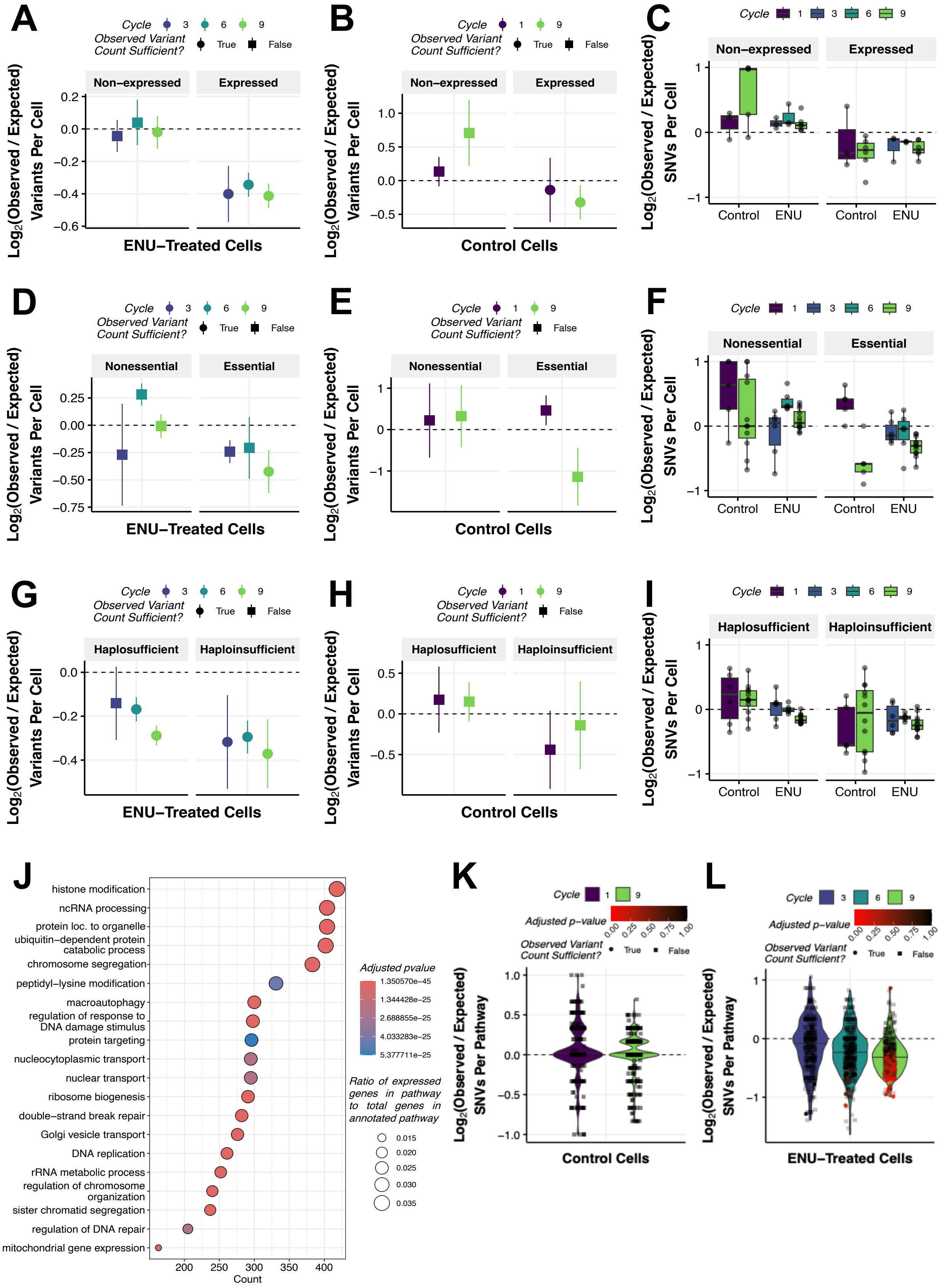

A
Cycle 3 6 9
Observed Variant Count Sufficient? True False
Non−expressed
Expressed
Log2(Observed / Expected) Variants Per Cell
ENU−Treated Cells
B
Cycle 1 9
Control Cells
C
Log2(Observed / Expected) SNVs Per Cell
Control
ENU
D
Nonessential
Essential
E
F
G
Haplosufficient
Haploinsufficient
H
I
J
histone modification
ncRNA processing
protein loc. to organelle
ubiquitin−dependent protein catabolic process
chromosome segregation
peptidyl−lysine modification
macroautophagy
regulation of response to DNA damage stimulus
protein targeting
nucleocytoplasmic transport
nuclear transport
ribosome biogenesis
double−strand break repair
Golgi vesicle transport
DNA replication
rRNA metabolic process
regulation of chromosome organization
sister chromatid segregation
regulation of DNA repair
mitochondrial gene expression
Count
Adjusted pvalue
Ratio of expressed genes in pathway to total genes in annotated pathway
K
Adjusted p−value
Log2(Observed / Expected) SNVs Per Pathway
L

**Supplementary Figure 6. Extended Selection Pressure Analysis of Mutation Accumulation at the Pathway Level. Related to Figure 6.**

**(A)** Unnormalized $Log_2$(O/E) ratio of SNVs in functional regions associated with genes determined to be non-expressed or expressed from bulk mRNA transcriptome data (Table S7). Related to Figure 6B. **(B)** Normalized $Log_2$(O/E) ratio of SNVs for control cells only in functional regions associated with genes determined to be non-expressed or expressed from bulk mRNA transcriptome data (Table S7). Related to Figure 6B. **(C)** Normalized $Log_2$(O/E) ratio of SNVs per cell in functional regions associated with genes determined to be non-expressed or expressed from bulk mRNA transcriptome data (Table S7). Related to Figure 6B. **(D)** Unnormalized $Log_2$(O/E) ratio of SNVs in functional regions associated with nonessential or essential gene sets expressed in bulk mRNA transcriptome data (Table S7). Related to Figure 6D. **(E)** Normalized $Log_2$(O/E) ratio of SNVs for control cells only in functional regions associated with nonessential or essential gene sets expressed in bulk mRNA transcriptome data (Table S7). Related to Figure 6D. **(F)** Normalized $Log_2$(O/E) ratio of SNVs per cell in functional regions associated with nonessential or essential gene sets expressed in bulk mRNA transcriptome data (Table S7). Related to Figure 6D. **(G)** Unnormalized $Log_2$(O/E) ratio of SNVs in functional regions associated with haplosufficient or haploinsufficient gene sets expressed in bulk mRNA transcriptome data (Table S7). Related to Figure 6F. **(H)** Normalized $Log_2$(O/E) ratio of SNVs for control cells only in functional regions associated with haplosufficient or haploinsufficient gene sets expressed in bulk mRNA transcriptome data (Table S7). Related to Figure 6F. **(I)** Normalized $Log_2$(O/E) ratio of SNVs per cell in functional regions associated with haplosufficient or haploinsufficient gene sets expressed in bulk mRNA transcriptome data (Table S7). Related to Figure 6F. **(J)** Top 20 gene ontology biological processes out of 399 resulting from gene set over-representation analysis of expressed genes from bulk mRNA transcriptome data from control and ENU-treated cells at cycle 9. Statistics: n=6; adjusted p-values estimated using a hypergeometric distribution test followed by Benjamini-Hochberg correction for multiple hypothesis testing. $Log_2$ O/E ratio of SNVs per cell in functional regions associated with genes determined to be non-expressed or expressed from bulk mRNA transcriptome data from control and ENU-treated cells at cycle 9. Related to Figures 6H-K. **(K)** Unnormalized $Log_2$(O/E) ratio of SNVs in the functional regions of expressed pathways for in bulk mRNA transcriptome data (Table S8). Related to figure 6H. Statistics: P-values for comparing the frequency of observed variants to that expected by chance alone were estimated using a permutation test (n=10,000) with Benjamini-Hochberg correction for multiple hypothesis testing. **(L)** Normalized $Log_2$(O/E) ratio of SNVs for control cells only in the functional regions of expressed pathways for in bulk mRNA transcriptome data (Table S8). Related to Figures 6H-K. Statistics: Same as in (K). Statistics for all panels: n=3 cells for cycles 1, 3, and 6; n=6 cells for cycle 9; Data represent mean ± S.D.

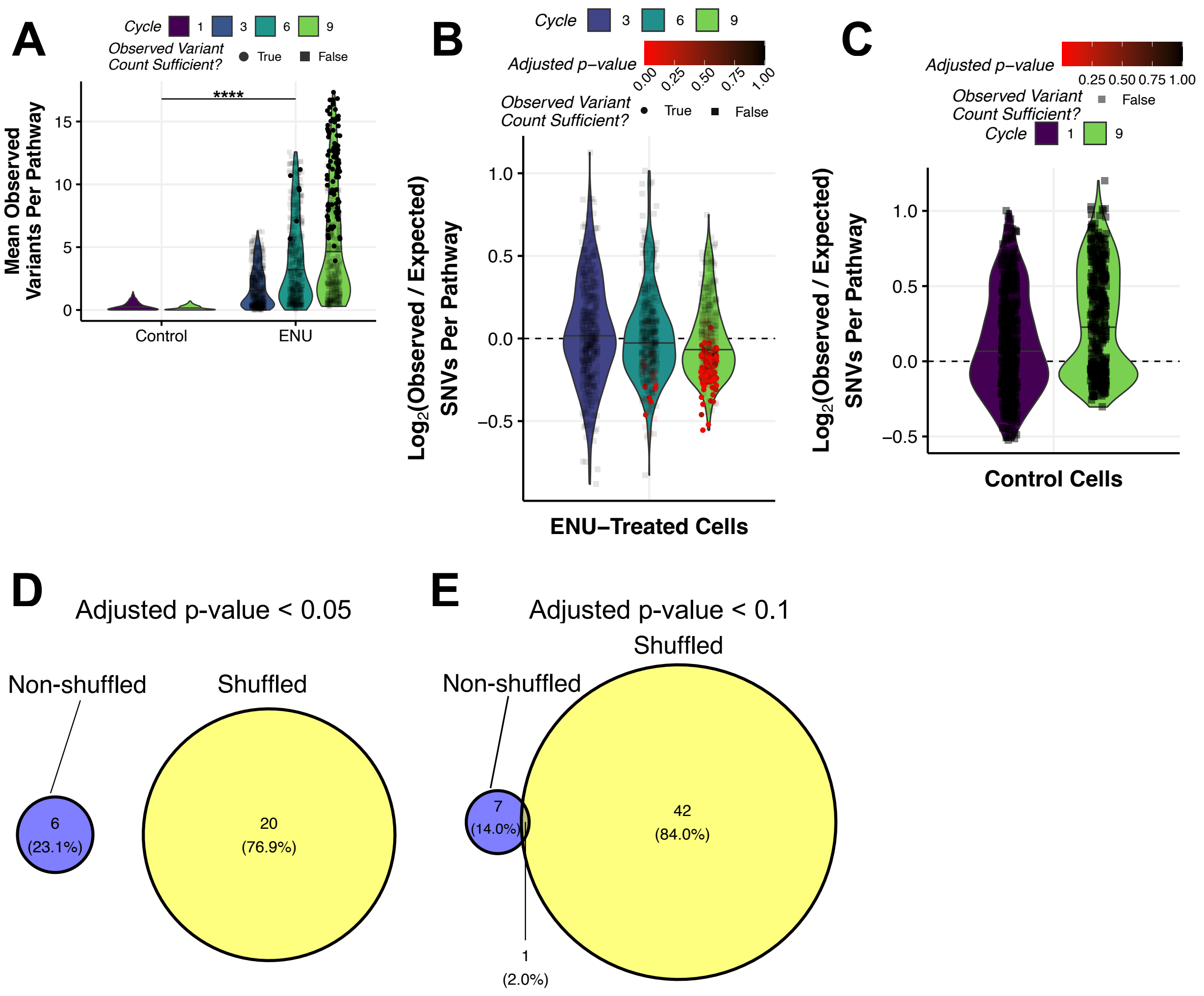

A
Cycle 1 3 6 9
Observed Variant Count Sufficient? True False
Mean Observed Variants Per Pathway
****
Control
ENU
B
Cycle 3 6 9
Adjusted p–value 0.00 0.25 0.50 0.75 1.00
Observed Variant Count Sufficient? True False
Log2(Observed / Expected) SNVs Per Pathway
ENU–Treated Cells
C
Adjusted p–value 0.25 0.50 0.75 1.00
Observed Variant Count Sufficient? False
Cycle 1 9
Log2(Observed / Expected) SNVs Per Pathway
Control Cells
D
Adjusted p-value < 0.05
Non-shuffled
Shuffled
6 (23.1%)
20 (76.9%)
E
Adjusted p-value < 0.1
Shuffled
Non-shuffled
7 (14.0%)
42 (84.0%)
1 (2.0%)

**Supplementary Figure 7. Extended Analysis of Negative Control to Validate Negative Selection of Functional Pathways. Related to Figure 6.**

**(A)** Mean observed frequency of SNVs within the functional regions of pathways expressed in bulk mRNA transcriptome data (Table S8) after shuffling genes between pathways (negative control). Statistics: n=399 pathways; p-values of comparisons of control and ENU-treated groups were estimated using a two-way ANOVA. ****: $p <= 0.0001$. **(B)** Mean $Log_2$(O/E) ratio of SNVs within the functional regions of expressed pathways expressed in bulk mRNA transcriptome data (Table S8) after shuffling genes between pathways (negative control). Statistics: P-values to assess if the $Log_2$(O/E) ratio significantly deviates from 0 were estimated using a permutation test (n=10,000) with the addition of Benjamini-Hochberg correction for multiple hypothesis testing of pathways. P-values were only estimated if the frequency of observed variants allowed for sufficiency statistical power (Figure S5B). **(C)** Normalized $Log_2$(O/E) ratio of SNVs for control cells only in the functional regions of pathways expressed in bulk mRNA transcriptome data (Table S8) after shuffling genes between pathways (negative control). Statistics: Same as in (B). **(D-E)** Overlap of pathways that show significant negative O/E ratios (i.e. pathways under negative selection) at cycle 9 in the ENU-treated group at an adjusted p-value<0.05 threshold (D) or an adjusted p-value<0.1 threshold (E) resulting from the original non-shuffled analysis (Figure 6H) or shuffled analysis (negative control; Figure S7B).

**Table S1. Sample table for bulk mRNA-sequencing experiment. Related to Figures 2 and S1.** Column headers: sample identifier (Sample), experimental condition (Condition), experimental cycle number (Cycle), biological replicate number (Replicate), and processing batch (Batch).

**Table S2. Differential gene expression results from bulk mRNA-sequencing experiment. Related to Figures 2 and S1.** Column headers: Ensembl gene ID (ensembl_gene_id), HGNC gene symbol (external_gene_name), Entrez gene ID (entrezgene_id), gene biotype (gene_biotype), chromosome (chromosome_name), start (start_position) and end (end_position) genomic coordinates, GC content (percentage_gene_gc_content), mean normalized expression (baseMean), log2 fold change between conditions (log2FoldChange), standard error of the log2 fold change (lfcSE), Wald test statistic (stat), raw p-value (pvalue), and Benjamini–Hochberg adjusted p-value (padj).

**Table S3. Sample table for single-cell whole genome sequencing experiment with metrics related to mutation calling and mutation burden. Related to Figures 3 and S2.** Column headers: Sample identifier (Sample), experimental condition (Condition), treatment cycle (Cycle), biological replicate number (Replicate), sequencing batch (Batch), total mapped bases at ≥20X coverage (Coverage (20X)), estimated sensitivity for SNV and INDEL detection (SNV sensitivity, INDEL sensitivity), observed SNVs and INDELs (Oberserved SNVs, Observed INDELs), estimated total SNVs and INDELs per cell (Estimated SNVs, Estimated INDELs), or per meagbase (Estimated SNVs per MB, Estimated INDELs per MB), and INDEL:SNV ratios (Observed INDEL:SNV ratio, Estimated INDEL:SNV ratio).

**Table S4. Results of selection analysis ($Log_2$(observed/expected)) for coding and non-coding variants. Related to Figures 5 and S4.** Column headers: Variant class (Variant Type), sample condition (Condition), treatment cycle (Cycle), observed variant count (Observed Variant Count), simulated mean of expected variants (Expected Variant Mean), normalized log2 observed/expected ratio (Normalized Log2(O/E)), standard deviation of the normalized log2 ratio (Normalized Log2(O/E) SD), required number of variants for statistical power (Observed Variants Required), whether this threshold was met (Observed Variant Requirement Met), unadjusted p-value (P-value), and FDR-adjusted p-value (Adjusted P-value).

**Table S5. Variant effect prediction annotation of all mutations.** See ensembl.org/info/genome/variation/prediction for information on columns.

**Table S6. Results of dndscv for cancer driver gene mutations in ENU-treated cells.** Column headers: gene name (gene_name), the counts of synonymous (n_syn), missense (n_mis), nonsense (n_non), splicing (n_spl), and indel (n_ind) mutations, the ratio or observed/expected metrics for each class (wmis_cv, wnon_cv, wspl_cv, wind_cv), the p-values for each mutation class (pmis_cv, pnon_cv, pspl_cv, pind_cv), combined measures for truncating or all substitutions (ptrunc_cv, pallsubs_cv), overall p-values for the gene (pglobal_cv), and the corresponding FDR-adjusted p-values (qmis_cv, qtrunc_cv, qallsubs_cv, qglobal_cv).

**Table S7. List of Entrez IDs for Non-expressed/Expressed, Nonessential/Essential, and Haplosufficient/Haploinsufficient gene sets. Related to Figures 6 and S5.** For Nonessential/Essential and Haplosufficient/Haploinsufficient gene sets, non-expressed genes were filtered out.

**Table S8. Gene set enrichment analysis results for gene ontology biological process of expressed genes. Related to Figures 6 and S5.** Column headers: GO term identifier (ID), associated biological process description (Description), ratio of input genes annotated to the GO term (GeneRatio), background gene ratio (BgRatio), raw p-value for enrichment (pvalue), multiple-testing adjusted p-value (qvalue), list of Entrez gene IDs contributing to the enrichment (Entrez gene ID), number of genes in the set (Count), and a custom enrichment score (qscore).

**Table S9. Results of selection analysis ($Log_2$(observed/expected)) for GO biological processes. Related to Figures 6 and S5.** Column headers: GO biological process term (GO biological process), GO term ID (GO ID), sample condition (Condition), treatment cycle (Cycle), number of observed variants (Observed Variant Count), simulated median of expected variants (Expected Variant Median), normalized log2 observed/expected ratio (Normalized Log2(O/E)), its standard deviation (Normalized Log2(O/E) SD), required variant count for statistical power (Observed Variants Required), whether this threshold was met (Observed Variant Requirement Met), raw p-value (P-value), and FDR-adjusted p-value (Adjusted P-value).

**Table S10. Catalogue of all mutations identified in control and ENU-treated cells. Related to Figures 3-6.** Column headers: Chromosome identifier (Chromosome), variant start and end positions (Start_Position, End_Position), reference allele (Reference_Allele), observed alleles in the sample (Seq_Allele1, Seq_Allele2), variant classification (Variant_Type), and sample identifier (Sample_ID).
